# A comprehensive macaque fMRI pipeline and hierarchical atlas

**DOI:** 10.1101/2020.08.05.237818

**Authors:** Benjamin Jung, Paul A. Taylor, Jakob Seidlitz, Caleb Sponheim, Pierce Perkins, Leslie G. Ungerleider, Daniel Glen, Adam Messinger

## Abstract

Functional neuroimaging research in the non-human primate (NHP) has been advancing at a remarkable rate. The increase in available data establishes a need for robust analysis pipelines designed for NHP neuroimaging and accompanying template spaces to standardize the localization of neuroimaging results. Our group recently developed the NIMH Macaque Template (NMT), a high-resolution population average anatomical template and associated neuroimaging resources, providing researchers with a standard space for macaque neuroimaging (Seidlitz, Sponheim et al., 2018). Here, we release NMT v2, which includes both symmetric and asymmetric templates in stereotaxic orientation, with improvements in spatial contrast, processing efficiency, and segmentation. We also introduce the Cortical Hierarchy Atlas of the Rhesus Macaque (CHARM), a hierarchical parcellation of the macaque cerebral cortex with varying degrees of detail. These tools have been integrated into the neuroimaging analysis software AFNI (Cox, 1996) to provide a comprehensive and robust pipeline for fMRI processing, visualization and analysis of NHP data. AFNI’s new @animal_warper program can be used to efficiently align anatomical scans to the NMT v2 space, and afni_proc.py integrates these results with full fMRI processing using macaque-specific parameters: from motion correction through regression modeling. Taken together, the NMT v2 and AFNI represent an all-in-one package for macaque functional neuroimaging analysis, as demonstrated with available demos for both task and resting state fMRI.

**Highlights:**

- The NMT v2, a stereotaxically aligned symmetric macaque template, is introduced.
- A new atlas (CHARM), defined on NMT v2, parcellates the cortex at six spatial scales.
- AFNI’s @animal_warper aligns and maps data between monkey anatomicals and templates.
- AFNI’s afni_proc.py facilitates monkey fMRI analysis with automated scripting and QC.
- Demos of macaque task and resting state fMRI analysis with these tools are provided.

## 1. Introduction

The macaque monkey brain serves as a useful model for understanding the organization and function of the primate brain. Rhesus macaques are phylogenetically close to and share many homologies with humans (Zhang and Shi, 1993), making the species an essential model for studying complex neural circuits and higher order cognition. Modes of investigating macaque neurophysiology involve noninvasive techniques, such as functional and structural neuroimaging (e.g. MRI), as well as invasive techniques including electrophysiology, lesions and chemogenetics. By combining modalities, researchers gain a better understanding of the nature of the primate nervous system.

While functional neuroimaging in nonhuman primates (NHPs) has revealed a great deal about the primate brain, researchers have historically been limited by the small amount of available data and the difficulty of acquiring new data. Macaque studies have generally relied on much smaller sample sizes than human studies, due to the difficulty of housing, training, and scanning animals. These challenges may be surmounted, and the statistical power of macaque neuroimaging studies could be strengthened, by multisite collaborations and data sharing. This was the motivation behind the development of the Primate Data Exchange (PRIME-DE; Milham et al., 2018) repository, which currently hosts NHP structural and functional neuroimaging data from 26 groups. These data are freely available to researchers, enabling larger multi-site studies. The PRIME-DE has been instrumental in leading this push towards openness and reproducibility in the macaque community (Milham et al., 2020), and data from the PRIME-DE has already been used in several studies (Froudist-Walsh et al., 2018; He et al., 2020; Heuer et al., 2019; Oligschläger et al., 2019; Valk et al., 2020; Xu et al., 2019b, 2019a, 2018).

The greater availability of macaque neuroimaging data is complicated by the variability in scanning protocols and the lack of established data analysis pipelines and group analysis tools. There is no single standard for neuroimaging data acquisition in the macaque, as evidenced by the PRIME-DE datasets, which vary with regard to scanning parameters such as field strength, voxel dimension and orientation, among a myriad of other differences (scanner-specific B0 inhomogeneities, coil sensitivities, etc.). Efforts are currently underway to provide standardized parameters for NHP data acquisition (Autio et al., 2020) and aggregate macaque neuroimaging techniques, pipelines and resources in the PRIMatE Resource Exchange (PRIME-RE; Messinger et al., this issue).

We observe that the research community would benefit from three areas of improvement in study design and implementation: 1) using anatomical templates to align and report results in standardized coordinate spaces; 2) having open, robust tools for the alignment of functional and anatomical data to these spaces (facilitating accuracy and visual verifications of spatial alignment); and 3) designing analysis pipelines that are robust, sharable and flexible for a wide array of study designs (so analyses can be understood and reproduced easily). Here, we present publicly available tools to address these three needs, demonstrating how they work both individually and synergistically.

Our group first addressed the issue of a common macaque MRI coordinate space in 2018 with the NIMH Macaque Template (NMT; Seidlitz, Sponheim et al., 2018). The NMT is a high-resolution template generated through iterative nonlinear alignment of 31 adult rhesus macaques; it was designed for group analyses and to facilitate collaboration and replicability in macaque research. The NMT has been used as a standardized space for a multitude of macaque imaging studies (Ainsworth et al., 2020; Arcaro et al., 2019; Ball and Seal, 2019; Bridge et al., 2019, 2018; Guehl et al., 2020; Menon et al., 2019; Milham et al., 2018; Ortiz-Rios et al., 2018; Seidlitz et al., 2018), and it serves as a standardized macaque space in which data can be aligned, displayed and analyzed within Neurovault (Gorgolewski et al., 2015), AFNI (Cox, 1996) and CIVET (Lepage et al., this issue).

The quality of the anatomical template can be a major limiting factor for alignment and, by extension, group analysis. Therefore, we introduce the latest updates to the NIMH Macaque Template, called NMT v2, which has improved contrast, various artifacts removed, and a symmetric representation of the macaque brain in stereotaxic coordinates. The NMT v2 is further augmented by the availability of atlases, which are critical for ROI-based analyses, especially in functional connectivity analyses. We describe a novel hierarchical atlas, the Cortical Hierarchy Atlas of the Rhesus Macaque (CHARM), that parcellates the NMT v2 cortical sheet at six different spatial scales. The addition of the CHARM to the NMT v2 allows researchers increased flexibility when choosing anatomical regions for analysis and provides parcellations of the macaque cortical sheet that are appropriate for characterizing the larger voxels typical of functional datasets. Both the NMT v2 and CHARM are available for download from the AFNI website.^1^ While there are many programs capable of performing fMRI analysis in macaques, most of these programs are designed for human MRI, and NHP neuroimaging can have different considerations and require different parameter settings. AFNI is a widely used program capable of performing a diverse set of functions related to neuroimaging processing and analysis that may be leveraged for NHP neuroimaging (Cox, 1996). We demonstrate AFNI’s capabilities with regard to NHP neuroimaging: including macaque-specific functional processing pipeline abilities and integration with the NMT v2.

As most group analyses require aligning data to a standard space, we first introduce a new registration tool, @animal_warper, which provides nonlinear alignment to the NMT v2 or any other anatomical scan. The program contains automatically generated quality control (QC) reports and applies the alignment to any additional atlases or segmentations provided.

Finally, we discuss creating flexible and reproducible analysis pipelines specifically for NHP neuroimaging using AFNI’s afni_proc.py program. We demonstrate @animal_warper and afni_proc.py with two macaque datasets (available from the AFNI website^2,3^), showing processing steps for resting state and task fMRI datasets. These start-to-finish processing steps include warping data to the NMT v2, and modeling responses if the contrast agent MION (monocrystalline iron oxide nanoparticle; Vanduffel et al., 2001) has been used. Taken together, the NMT v2 and AFNI provide a comprehensive and flexible analysis pipeline for macaque fMRI data.

## 2. Materials and Methods

### 2.1 Subject information

The NMT v2 was created from T1-weighted scans of 31 rhesus macaques (*Macaca mulatta*) from the Central Animal Facility at the NIMH, as described in (Seidlitz, Sponheim et al., 2018). An additional 10 macaques were scanned in a horizontal 3T scanner at the NIH Clinical Center for calculating how to place the NMT v2 in its stereotaxic orientation. All animal procedures were conducted in compliance with the National Institutes of Health Guide for the Care and Use of Laboratory Animals.

This project additionally uses openly available PRIME-DE^4^ datasets from the National Institute of Mental Health - Messinger (NIMH), the Netherlands Institute for Neuroscience (NIN) and the Stem Cell and Brain Research Institute (SBRI), the details of which can be found in Milham et al. (2018).

### 2.2 The NIMH Macaque Template version 2 (NMT v2)

The NMT v2 templates were created through iterative nonlinear alignment of 31 rhesus macaque scans. Compared to the original NMT (hereafter referred to as NMT v1.2), the NMT v2 templates have improved template contrast and several artifacts have been removed. To facilitate surgical planning and reporting of coordinates in a widely-used coordinate system, the NMT v2 was reoriented to approximate the brain’s orientation in a Horsley-Clarke stereotaxic apparatus (Horsley and Clarke, 1908). Brain masks and segmentation were manually refined for each NMT v2 template, and we provide templates in multiple fields of view and resolutions. The remainder of this section describes the making of the NMT v2 in greater detail.

#### 2.2.1 Generation of the NMT v2 Templates

NMT v2 templates were generated using a modified version of the previously described NMT creation pipeline (Seidlitz, Sponheim et al., 2018). The NMT v2 templates were created through 4 iterations of nonlinear alignment to an intermediate registration target and averaging of the aligned scans. The NMT v2 template creation pipeline differs from the NMT v1.2 template creation at the final averaging step. After the final iteration of nonlinear registration, the average at each voxel was restricted to values between the 20^th^ and 80^th^ percentiles of voxel intensities across the 31 volumes to prevent outliers from skewing this group average. This averaging procedure was used to reduce or eliminate several artifacts present in the NMT v1.2 (see section 3.1.1). The final voxel resolution of the template matched the resolution of the up-sampled input scans (0.25 mm isotropic voxels).

The symmetric NMT v2 template was created using a similar process, except each anatomical scan was first duplicated and mirrored about the plane between the hemispheres. The resulting 62 volumes were then nonlinearly aligned to an intermediate template over 4 iterations. To remove the minor remaining asymmetries, the right hemisphere was reflected and replaced the left hemisphere to create a perfectly symmetric template. No information was lost during this process, as both hemispheres contained the same input data from 62 hemispheres across 31 subjects.

#### 2.2.2 Reorientation of the NMT to the Horsley-Clarke Stereotaxic Apparatus

The NMT v1.2 was defined using a standardized space with the origin at the center of the anterior commissure, and the horizontal plane passing through the superior extent of the anterior commissure (AC) and the inferior extent of the posterior commissure (PC) (essentially AC-PC alignment) (Reveley et al., 2017). Surgical procedures, electrophysiological recording, drug infusions and other techniques require precise brain coordinates and angles of approach. To obtain such precise and reproducible coordinates, an anesthetized macaque would typically be placed in a Horsley-Clarke stereotaxic apparatus, in which the horizontal plane is defined by the center of the interaural meatus (ear bars) and the infraorbital ridge (Horsley and Clarke, 1908). Collecting scans of the monkey in an MRI-compatible stereotaxic apparatus prior to surgery allows researchers to visualize particular brain structures, determine their precise location, fabricate any custom apparatus, and chart out a surgical approach.

To get the NMT v2 into stereotaxic orientation, we collected anatomical scans (T1-weighted) from 10 anesthetized rhesus macaques, while they were in an MRI-compatible Horsley-Clarke stereotaxic apparatus. An affine transformation was used to register the AC-PC aligned NMT v1.2 to each of these stereotaxic scans, but only the pitch rotation value was extracted from the resulting affine matrices.^5^ The average pitch angle was then used to reorient the NMT into stereotaxic space. With this process we were also able to assess the consistency of stereotaxic placement by calculating the variance of the pitch angle within subjects placed in a stereotaxic apparatus on different sessions (discussed further in section 3.1.2). The origin of the NMT v2 coordinate system was placed at the intersection of the midsagittal plane and the interaural line.

#### 2.2.3 Post-processing

Voxel intensities of the templates were capped based on the distribution of blood vasculature intensities. Capping the upper range of values helps make the other, darker tissue types easier to differentiate in a standard neuroimaging viewer.

Several maps of the NMT brain tissue, including a brain mask, tissue segmentation, and a mask of the cerebellum, were previously generated for the NMT v1.2 in Seidlitz, Sponheim et al. (2018). Despite differences in morphology between the NMT v1.2 and the NMT v2 templates, nonlinear alignment could be used to accurately map these structures in the new templates. We calculated the nonlinear alignment between the NMT v1.2 and the NMT v2 templates using AFNI’s 3dQwarp (Cox and Glen, 2013), and the NMT v1.2 masks were brought to the NMT v2 using 3dNwarpApply with nearest neighbor interpolation. The masks were then manually refined in the symmetric and asymmetric NMT v2 templates to better match the NMT v2 morphology. Masks for the symmetric NMT v2 template were additionally symmetrized by mirroring the values from the right hemisphere.

Bias field correction was performed on the NMT v2 templates using the N4 algorithm (Tustison et al., 2010). The b-spline was fit only to white matter (WM) voxels, defined using the warped segmentation of the NMT v1.2, which we observed to provide better bias field correction compared to a N4 bias field correction performed within the whole brain mask. The bias field was extrapolated and applied across the entire volume, thereby eliminating artifacts at the brain mask boundary (between bias-corrected and non-bias corrected voxels) present in the NMT v1.2. Following N4 bias field correction, voxel intensities were again capped to limit the distribution of values and then scaled to a maximum value of 1024 (arbitrary units). All volumes were converted from “float” to “short” data type to save space and improve processing efficiency.

The grid of the NMT v1.2 had an odd number of voxels in the x-direction. With the introduction of the symmetric NMT v2, we switched to a grid with an equal number of voxels in the left and right hemispheres to simplify interhemispheric comparisons. Consequently, the midline of the NMT v2 corresponds to a voxel edge as opposed to the middle of a voxel (a 0.125 mm shift).

#### 2.2.4 NMT v1.3: back-applying methodological improvements to NMT v1.2

As highlighted above, several of the processing steps applied in creating NMT v2 differed from the original NMT v1.2 creation, and these appeared to improve local features (tissue contrast, etc.) without changing the underlying morphology of the template. We therefore regenerated the NMT v1.2 and applied these morphology-preserving steps, resulting in the NMT v1.3.

In this updated version, the previously described N4 bias field correction and “short” scaling were applied to the regenerated NMT v1.2 template. We maintain the same FOV and grid dimensions as the NMT v1.2, and therefore the NMT v1.3 has odd matrix dimensions. We also include the CHARM in this NMT v1.3. (see section 2.3 for details). The NMT v1.3 is currently available on Github^6^ and the AFNI website^7^. Users should expect to be able to use the NMT v1.3 in place of NMT v1.2 with only minimal differences. All volumes associated with the NMT v1.3 were assigned a standard *NMT* space label via the AFNI extension in the NIFTI header.

#### 2.2.5 NMT v2 Segmentation

The NMT v2 templates have been segmented into 5 tissue classes: cerebrospinal fluid (CSF), WM, subcortical gray matter (scGM), cortical gray matter (GM) and arterial blood vasculature (BV). WM, GM and CSF were defined primarily by segmenting the NMT v2 templates using the new CIVET-macaque pipeline (Lepage et al., this issue). The cerebellum and scGM, which are often not well delineated by automated segmentation algorithms, were manually defined. The scGM outside the cerebellum was defined using regions in the new Subcortical Atlas of the Rhesus Macaque (SARM; Hartig et al., this issue), which were manually defined based on the NMT v2 morphology. BV was defined using a combination of high intensity values and blood vasculature maps from the CIVET-macaque pipeline. Minor errors in tissue classification were manually refined in ITK-SNAP (Yushkevich et al., 2016) and AFNI.

#### 2.2.6 NMT v2 Field of View

The field of view (FOV) of anatomical scans can vary widely based on acquisition strategies. Some are restricted almost exclusively to the brain, whereas others may include a large amount of the surrounding head, neck and shoulders and may not be centered on the brain. Additionally, scans may be collected with the macaque in different positions, such as the sphinx position or a seated position in a vertical scanner. These differences in FOV and positioning can interrupt alignment algorithms, resulting in poor alignment, and/or require additional manual processing interventions. The default NMT v2 FOV covers the brain and some surrounding CSF. We have observed that alignment of scans with FOVs much larger than this tends to be better when the template also has an expanded FOV. Therefore, the NMT v2 is distributed with two FOVs: the “standard” FOV (slightly larger than the brain; default) and a “full head” FOV (captures the entire macaque skull). The standard FOV is likely preferable for most uses of the NMT, including single subject alignment and segmentation, because the limited FOV reduces processing time and resources compared to the “full head” NMT.

#### 2.2.7 NMT v2 Spatial Resolution

The native voxel resolution of the NMT v2 is 0.25 mm isotropic. This spatial resolution captures the most detail about the macaque brain, reduces partial volume effects and achieves accurate registration based on detailed features. However, in practice, macaque anatomical scans are rarely collected at resolutions much finer than 0.5 mm isotropic (and most functional volumes are much coarser). While higher resolution templates are useful for some analyses, in many cases lower resolution base volumes would provide essentially the same outputs, with the benefit of lower processing times and disk space. Therefore, for convenience, we also provide the NMT v2 datasets at 0.5 mm isotropic resolution. We note that in human fMRI studies, EPI voxel edges are typically 2-3 mm and those of anatomical volumes/templates are 1 mm; in macaque imaging, where EPI voxel edges are typically 1.5-2 mm, having anatomical volumes and templates with voxels of 0.5 mm preserves a similar size ratio.

#### 2.2.8 The NMT2 Standardized Coordinate Space

Since the release of the NMT v1.2, NIFTI standard format headers (Cox et al., 2004) have expanded the specifications for *qform* and *sform* to allow for the specification of normalized coordinate spaces beyond the human Talairach and MNI spaces (*[qs]form_code* = 5). We therefore have defined a new standardized space designation, *NMT2*, to represent our new NMT v2 templates in stereotaxic coordinates. The *NMT2* space is compatible with AFNI’s *whereami* feature, allowing users to associate *NMT2* coordinates with specific atlas structures either in the GUI or via the command line for any dataset aligned to the NMT v2.

### 2.3 The Cortical Hierarchy Atlas of the Rhesus Macaque (CHARM)

The Cortical Hierarchy Atlas of the Rhesus Macaque (CHARM) is a novel six-level anatomical parcellation of the macaque cerebral cortex. The levels provide a progressively finer parcellation of the cortical sheet, permitting users to select regions at a range of spatial scales. The variety of organizational spatial scales in the CHARM make the atlas suitable for a wide array of functions and analyses. Broad-scale levels are useful for indicating general regions, such as those with names conserved across several mammalian species, and for observing large-scale activations or conducting cross-species analyses. Fine-scale levels are useful for differentiating between cytoarchitectonic subdivisions of a cortical area or specifying the target of a local injection or neuronal recording. Intermediate-scale levels are well suited for selecting a complete cortical area (e.g. a Brodmann area) or a group of areas that are either functionally similar or in close proximity (e.g. areas on the same gyrus). Different scales may also be used in tandem. For example, a seed region can be described at a fine scale and its resting state functional connectivity can be described more broadly. The same logic may also be used to track the projections of a tracer injection or the influence of a brain perturbation technique. This flexibility is important for distilling whole brain neuroimaging data in ways that respect the scale of the results. The choice of CHARM level also allows the user to determine *a priori* how many regions will be employed in an analysis and thus the required degree of multiple comparison correction.

The CHARM hierarchy starts with broad structures that are progressively subdivided. The first and broadest level of CHARM divides cortex into the frontal, parietal, temporal, and occipital lobes. In the second level, each lobe is divided into 3-6 large regions, with common names that are frequently also used in humans and some other mammalian species (e.g. motor cortex, superior parietal lobule [SPL], medial temporal lobe [MTL]). In the third through sixth levels of CHARM, regions from the preceding level are typically subdivided into 2-3 smaller structures (up to 4). Some regions remained unchanged between one or more successive levels. Levels 2 and 3 mostly consist of fairly large cortical divisions (e.g. lobules, surfaces of a lobe or gyrus) or groups of related areas (e.g. temporal pole, core areas of auditory cortex, lateral motor cortex [M1/PM]). Level 4 and 5 regions are typically composites of a few related cortical areas (e.g. Brodmann areas) or larger subdivisions of a single cortical area. Level 6 areas are mostly fine subdivisions of cortical areas (or the whole area if it has not been subdivided). This finest level matches the parcellation of the D99 digital atlas (Reveley et al., 2017), except for some changes in nomenclature and the fusion of the smallest D99 structures with one or more neighbors. Sections <u>2.3.1</u> and <u>2.3.2</u> describe the creation of the CHARM.

#### 2.3.1 Segmentation-based refinement of the D99 Atlas on the NMT v2

The first step in the creation of the CHARM was to accurately map the D99 macaque atlas from the D99 template to the NMT v2 templates. During visual inspection, we found discontinuities in two regions (areas 1-2 and 24a) of the D99 atlas v1.2a and manually repaired these discontinuities using the Saleem and Logothetis (2012) atlas as a guide. The D99 template was then nonlinearly warped to the NMT v2, and this transformation was applied to the corrected D99 atlas. The transformed atlas covered much of the NMT v2 but left some cortical voxels unlabeled. In addition, there were several places where a cortical region from one sulcal bank spread across both banks of the sulcus after the warping procedure.

The D99 is an *ex vivo* template so the opposing banks of sulci are frequently abutting, whereas the NMT v2 templates are based on *in vivo* scans, and so sulcal banks are separated by CSF. To improve the assignment of labels to appropriate voxels, we used a segmentation-based refinement method. The cortical D99 regions were masked using the cortical GM segmentation of the NMT v2 templates, thereby removing atlas labels from the CSF and splitting anatomical regions that had spread across the sulcal banks. We preserved the largest contiguous region and removed the smaller erroneous clusters. We then dilated each region using AFNI’s 3dROIMaker (Taylor and Saad, 2013) to fill in any voxels within the cortical sheet that were left unlabeled during the registration process or that had their labels removed during the masking and clustering process. Finally, the subcortical regions, relatively unaffected by the transition from *ex vivo* to *in vivo* morphology, were combined with the modified cortical regions to recreate the D99 atlas in the NMT v2 templates. The cortical regions of this refined D99 atlas were used as the starting point for making the CHARM.

#### 2.3.2 Defining the levels of the CHARM

Each level of the hierarchical atlas partitions the full cortical sheet, including neocortex and allocortex, by successively combining regions to form composite regions. At every level, the CHARM borders follow the borders of the D99 atlas cortical regions warped to the NMT v2. At the finest level of CHARM (level 6), most regions are identical to the individual D99 regions, except that regions smaller than 13.5 mm^3^ in the D99 atlas (equivalent to the volume of 4 isotropic voxels of side 1.5 mm) were combined with one or more neighboring regions. Due to their small size and geometry, several (especially allocortical) D99 areas are not consistently present or contiguous after registering this 0.25 mm isotropic atlas to 1.5 mm isotropic functional scans, (see section 4.2). Fusing the smallest D99 regions with a neighbor produced CHARM regions large enough to survive nonlinear warping to individual subjects and down-sampling to fMRI resolutions for use in ROI based analyses.

We defined broader levels of the CHARM by combining adjacent regions to form larger composite structures. Each region belongs to only one composite structure in each broader level so that collectively all the areas in a given level represent the cortex once and only once. Regions were combined based on various considerations including geographical proximity and similarity in terms of cytoarchitecture, function, nomenclature, and anatomical connectivity. Initially, subdivisions were combined to form complete cortical areas (e.g. areas 45a and 45b were combined to form area 45), with intermediate groupings sometimes used to form larger subdivisions of the cortical area (e.g. the 6 subdivisions of the temporal pole were first combined to make granular, dysgranular, and agranular subdivisions). Cortical areas were then combined on the basis of spatial and/or functional characteristics (e.g. areas 32, 25, and 24 were combined to form the anterior cingulate cortex; area 3a/b was combined with areas 1-2 to form primary somatosensory cortex). These collections of cortical area were then fused to form progressively larger ensembles (e.g. dorsolateral prefrontal cortex, posterior cingulate gyrus, hippocampus, belt areas of auditory cortex). At level 1, the cortex was divided into the four lobes. For examples of the CHARM organization, refer to section 3.2.1.

Where possible, the CHARM nomenclature follows existing names for larger cortical groupings. At the finer levels, the CHARM names and abbreviations for areas largely follow the D99 atlas. One exception is in the motor cortex (see Table 2.2 of Saleem and Logothetis (2012) for equivalencies). Another is that CHARM area names generally do not refer to composite structures provided in another CHARM level. So, for example, the D99 area “subregion [24a] of anterior cingulate cortex” is called just “area 24a” in CHARM level 6, as its affiliation with the anterior cingulate cortex is made explicit in CHARM level 3. Each CHARM region has both a unique name and abbreviation. For purposes of programming compatibility, the first character of these abbreviations are not numbers, spaces have been replaced with underscores, and reserved special characters (e.g., “?”) are not used.

While there are a myriad of ways to group cortical regions, our choices were guided by well-established larger cortical divisions and previous work including both paper and digital macaque brain atlases (Saleem and Logothetis, 2012, especially chapter 2; Bowden and Martin, 2000; Paxinos et al., 2008; Rohlfing et al., 2012), digital resources^8^ (Dubach and Bowden, 2009), anatomy studies (Kravitz et al., 2013, 2011; Scott et al., 2017), neuroinformatic approaches (Caminiti et al., 2017), and related hierarchies of monkeys^9^ and humans^10^.

There were exceptions to and conflicts between many of the grouping principles described above. While all CHARM regions were confined to a single cortical lobe, they were not all contiguous. Areas V3 and V4 are composites of their discontinuous dorsal and ventral subdivisions. These exceptions were made on functional grounds because together the dorsal and ventral subdivisions form complete maps of the visual field. There were also exceptions to subdivisions combining to form a single cortical area. There is no area 7 in the CHARM because its medial subdivision is not contiguous with the area 7 subdivisions in the inferior parietal lobule. Similarly, the CHARM does not include an area 8, 12, or 46 because the subdivisions of these areas are situated on different portions of the cortical surface. For example, in the case of area 12, the medial and orbital subdivisions (level 5) are located in the orbital frontal cortex (level 2), whereas its rostral and lateral subdivisions (level 5) are part of the lateral prefrontal cortex (level 2). Similarly, the lateral and ventromedial intraparietal sulcus regions (level 3) do not fuse because they belong to the superior and inferior parietal lobule, respectively (level 2). Likewise, the core and belt areas of the auditory cortex do not fuse with the parabelt areas of the auditory cortex because the latter belongs to the caudal superior temporal gyrus.

### 2.4 AFNI’s @animal_warper: nonlinear alignment to template space

Here we introduce @animal_warper, a new AFNI program for alignment of one animal’s MRI with another MRI, typically a species-specific template. The alignment process produces not only the subject anatomy registered to the space of the template, but also the affine (approximate, but quick) and nonlinear (full, but more computationally intensive) transformations between the spaces. Using these transformations, @animal_warper can also bidirectionally map any accompanying datasets (e.g., fMRI data, ROI maps or masks) between the native and the template spaces. For example, @animal_warper can transform the template’s brain mask to the native space for purposes of template-based skull stripping. Similarly, an atlas in the standard space can be transformed to the subject’s anatomical, providing a parcellation of the subject’s brain in its native space. Alternatively, maps in the native space, such as functional activations or other anatomical datasets (e.g., T2-weighted or FLAIR volumes), can be transformed to a standard space for comparison of subjects in a common template or to create a probabilistic map.

The user inputs a subject’s anatomical (e.g., T1-weighted) “source” dataset and a reference “base” volume to be aligned to (e.g., NMT v2). Optionally, an atlas or tissue segmentation map in the template space can also be included for mapping to the subject’s native space. The program works by first capping outlier voxel intensities in the volume, then aligning the source center to that of the template, followed by affine and nonlinear alignment. The optional atlas and/or segmentation datasets are warped into the original space using the inverse warps. Skull stripping of the subject’s anatomical is provided by masking with the skull stripped template or a template brain mask. Quality control images of the alignment and transformation of atlas and segmentation datasets are also created as montages in multiple planes (i.e. axial, sagittal, and coronal views). Surfaces are made for all the mapped atlas regions, as well as for the mapped template surface, in native space; scripts to open and view these surfaces in the SUMA GUI (Saad et al., 2004) are also automatically created. Additionally, summary “report” tables are generated for each atlas and segmentation, describing the volumes of ROIs before and after warping, as well as their fractional volumes and relative volumes.

The processing of macaque and other non-human species’ MRI data presents additional challenges that are different from the longer studied human-based MRI pipelines. For example, the size scale of anatomical structures is different, and this can require parameter adjustment for spatial processing, such as alignment. As such, one can set the “feature_size” in millimeters to specify the minimal spatial scale to try to match during alignment (both in @animal_warper and in some other AFNI programs); for most macaque MRI datasets, we have generally used 0.5mm as the feature size. Blurring and other spatial features that depend on size can be tuned to the experiment and analysis, varying with species and imaging type. For example, in ROI-based analyses, one might exclude blurring in favor of averaging the signal within a given ROI.

It should be noted that @animal_warper makes use of AFNI’s main nonlinear alignment program, 3dQwarp, in conjunction with other programs. While nonlinear warping is in general computationally intensive, 3dQwarp---and hence, a major component of @animal_warper---is inherently parallelized using OpenMP (www.openmp.org) and therefore can take advantage of computing systems that contain multiple CPUs to reduce overall runtime. Users can control the number of CPUs used by @animal_warper (subject to system constraints), and examples of doing so in processing scripts are provided in each of the AFNI macaque demos. For the 6 anatomicals processed across the AFNI demos, the mean runtime of @animal_warper on a laptop using 6 CPUs with 1.80 GHz processors was 30.3 minutes (min/max = 22.0/51.0).

Nearest neighbor interpolation is typically used to transform ROI labels from source to a target grid. In that method, each voxel is assigned the label of the closest voxel from the transformation from the source grid. Nonlinear transformations can twist these regions into non-plausible shapes in the target, often producing spatial discontinuities. We introduce an option in @animal_warper called “modal smoothing” that can act as a method of regularization. This procedure recomputes the nonlinearly warped voxel labels based on the most common label value (i.e. the mode) in a local neighborhood around every voxel. This method enforces at least local contiguity within a spherical neighborhood (default: 1 voxel radius), which results in more realistic looking ROI boundaries. Modal smoothing may be performed on datasets outside of the @animal_warper pipeline using AFNI’s 3dLocalstat program.

### 2.5 AFNI’s afni_proc.py: creating macaque fMRI processing streams

AFNI’s afni_proc.py is a well-established program for generating complete fMRI processing pipelines for individual subjects in both human and nonhuman imaging. The program takes a hierarchical approach to designing pipelines: first, the researcher specifies major processing “blocks” to be undertaken (e.g., time series despiking, various alignments, and regression modeling) and provides any necessary inputs (e.g., anatomical and EPI datasets, other datasets such as tissue maps or parcellations, stimulus timing files, and physiological measurements); then, options can be provided to control the implementation of each processing block (e.g., blur radius, hemodynamic response function, censoring thresholds). Using these processing blocks, the program generates a fully commented script that utilizes AFNI commands to process the given data. In this way, the researcher can generate fully customized processing pipelines from a compact afni_proc.py command (typically, 10-40 options are provided).

The hierarchical nature of afni_proc.py means that processing streams can be easily updated or adjusted (e.g. adding or changing options, removing runs with poor behavior or excessive movement), and re-run. There is no need to manually alter the flow of a complicated processing script and/or to carefully propagate small changes through many lines of code. The afni_proc.py command is succinct enough to be published, for example in a paper’s appendix, and can be easily shared, both with other groups and within a lab: afni_proc.py’s processing streams are inherently reproducible, because the pipeline can be recreated by knowing the command, which is saved in the output script, and the version of AFNI used.

The afni_proc.py command also automatically generates QC features. Both quantitative features (e.g., temporal signal-to-noise ratio (TSNR), degree of freedom counts, censor-based warnings and relative left-right consistency (Glen et al., 2020)) and qualitative features (e.g., images to show alignment and statistical maps) are produced and stored in a single, navigable HTML file for efficient quality control by the user. These QC reports provide a convenient and systematic way to review all the steps of processing, rather than just glancing at end results. Such reviews are especially important in NHP imaging, with its many idiosyncrasies.

Historically, afni_proc.py has primarily developed with a view toward human studies, but its generalized framework and flexibility of specification means that it is easily implemented on macaques (and other species). Indeed, afni_proc.py has macaque-tailored features, such as the option to select a hemodynamic response function appropriate for MION contrast. Furthermore, macaque-specific templates, such as the NMT v2, can be used as a standard space, and the nonlinear warps estimated by @animal_warper can be input directly into afni_proc.py and concatenated with other alignments (i.e. from anatomical alignment and registration of EPI volumes across time) to avoid repeated interpolation from multiple re-gridding processes. New features are constantly being added to AFNI by the developers in response to users’ requests and so AFNI will continue to adapt in response to the needs of animal studies and researchers.

### 2.6 fMRI processing demos

There are currently two fully interactive fMRI processing demos for macaque datasets in AFNI (a task fMRI demo and a multi-subject resting state fMRI demo), each containing basic datasets, commented processing scripts and a README.txt description file. We briefly summarize each demo here; for further details, see the demo processing scripts (as well as the PRIME-DE website for dataset-specific details).

The MACAQUE_DEMO_2.0 demo contains task fMRI data (TR=3000 ms, voxel size = 1.5 mm isotropic) plus a typical anatomical volume (voxel size = 0.5 mm isotropic) from a single macaque. During the EPI acquisition, the subject was shown images from one of four categories (face, object, scrambled face, scrambled object) in 36 s blocks. There are 13 fMRI runs of 112 volumes each, and MION contrast was present. Briefly, the anatomical dataset was first aligned to the “low-res” symmetric NMT v2 with @animal_warper. Then afni_proc.py was used to setup a standard processing pipeline, including: motion estimation, with censor limits at 0.2 mm and 2% outlier fraction in mask; blurring with a 2 mm radius; the application of @animal_warper linear and nonlinear transforms to bring EPI data into standard NMT v2 space; block design modeling the hemodynamic response function (HRF) with MION contrast agent over a 36 second block (“MIONN(36)” in afni_proc.py) and the inclusion of several general linear tests (GLTs) based on the estimated stimuli betas, such as “face vs scrambled face,” “face vs object,” etc. This pipeline is depicted schematically in Figure 1.

**Figure 1.**
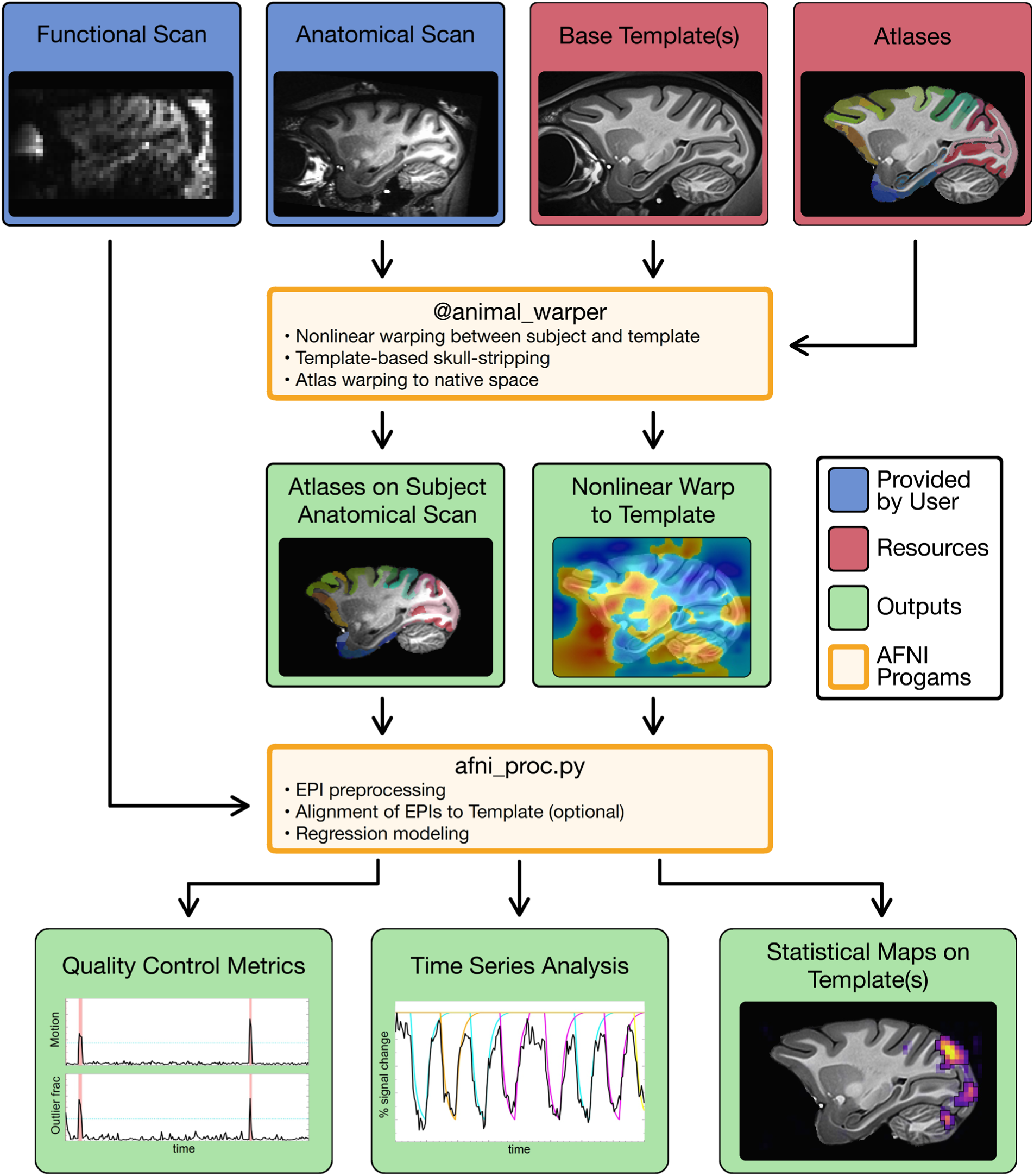
An example overview of task-based fMRI analysis in AFNI. Unprocessed functional and anatomical scans are collected by researchers in a native subject space (top row, blue). For voxel-wise analyses, the fMRI datasets are typically aligned to a common space. AFNI’s @animal_warper program estimates nonlinear alignment between a subject’s anatomical and any target template (e.g., the symmetric NMT v2; top row, red). The calculated warp field (middle row, right, green) is applied to align atlases, segmentations, and other maps in the template space (here the CHARM level 5; top row, red) to the subject (middle row, left, green) and to align native space volumes to the template. The nonlinear warps are saved for future applications. Images and tables of ROI information are automatically created for efficient QC purposes. The native space anatomical and functional datasets, in combination with @animal_warper’s nonlinear transforms, and any supplementary files of experimental design, such as stimulus timings for task fMRI, may then be supplied to afni_proc.py. afni_proc.py generates complete single subject processing streams that perform motion correction and other preprocessing steps, apply the nonlinear warps to bring all functional data to the NMT v2, and perform regression modeling. QC metrics and images are created for rapid exploration of all steps and final outputs (bottom row, green), including motion and outlier profiles, models of hemodynamic response for task fMRI, and volumetric statistical maps.

The MACAQUE_DEMO_REST demo contains datasets from 6 PRIME-DE macaques: 3 awake from NIMH (Messinger et al.; one subject is the subject analyzed in the MACAQUE_DEMO_2.0, with the same anatomical volume but different EPI data), 2 anesthetized from NIN (Klink and Roelfsema), and 1 anesthetized from SBRI (Procyk, Wilson and Amiez). For each subject, we include two resting state EPI scans and one anatomical T1-weighted anatomical volume. Across the datasets, EPI voxel edge lengths range from 1.25 to 1.70 mm, repetition time (TR) values range from 1700 to 3000 ms, and run lengths range from 200 to 420 volumes; anatomical resolution is either 0.5 or 0.6 mm isotropic. As in the task fMRI demo, @animal_warper was first run on each subject’s anatomical data, generating warps to the “low-res” symmetric NMT v2. Here, both awake and anesthetized data were processed as standard resting state acquisitions, including: motion correction, with both motion estimates and their derivatives; motion censoring at a limit of 0.2 mm and outlier fraction of 5% per volume; and inclusion of nonlinear warping via @animal_warper. For voxel-wise analyses, one might blur with a 2 mm radius; for ROI-based analyses, blurring would not be included.

### 2.7 Data accessibility and availability

The NMT v1.2, NMT v1.3, NMT v2 templates and the CHARM are all freely distributed via AFNI and compatible with any program using the NIFTI/GIFTI format. Templates and atlases may be downloaded via the AFNI website^11^, as well as with the AFNI command @Install_NMT. The @animal_warper.py and afni_proc.py programs are distributed within the open source program AFNI (Cox, 1996). All these resources are also available via the PRIME-RE portal (Messinger et al., this issue). AFNI binaries are downloadable from the AFNI website^12^, and the source code is available on Github^13^.

The complete demo examples (including both data and scripts) for processing the macaque functional data can be downloaded with the AFNI commands @Install_MACAQUE_DEMO (task fMRI demo) and @Install_MACAQUE_DEMO_REST (resting state fMRI demo). More information on these demos may be found in the AFNI online documentation^14,15^.

The anatomical scans used to generate the NMT v2 have also been processed using the CIVET-macaque pipeline to produce surface maps of each individual (Lepage et al., this issue). These will be released on the PRIME-DE in the near future.

## 3. Results

### 3.1 The NIMH Macaque Template v2

In the version 2 release of the NIMH Macaque Template (NMT v2), we provide two anatomical templates of the macaque brain in a Horsley-Clarke stereotaxic orientation (Horsley and Clarke, 1908): a symmetric template useful for general studies and interhemispheric comparisons, and an asymmetric template that preserves population-level differences between the two hemispheres. The symmetric and asymmetric variants of the NMT v2 are morphologically similar (Figure S1). The standard form of each template has 0.25 mm isotropic resolution and a FOV that covers the entire brain. For particular applications, we also provide each template at a lower resolution and in a larger FOV. The “low-res” NMT v2 has a reduced 0.50 mm isotropic resolution and the “full-head” NMT v2 has an expanded FOV that covers the entire skull. All variants of the symmetric and asymmetric template come with a brain mask, tissue segmentation, as well as the CHARM, the SARM and the D99 atlases. We also provide surfaces of the NMT v2 templates, generated through the CIVET-macaque pipeline (Lepage et al., this issue). The NMT v2 templates provide a flexible space for macaque neuroimaging with integrated resources for masking your data and conducting ROI-based analyses.

#### 3.1.1 Comparison to NMT v1.2

A summary of the similarities and differences between the original NMT v1.2 (Seidlitz, Sponheim et al., 2018) and the NMT v2 is provided in Table 1. The NMT v1.2 is an asymmetric-only template positioned in an AC-PC horizontal alignment with its origin in the AC. In contrast, the NMT v2, which has both a symmetric template and an asymmetric template, is in stereotaxic orientation with its origin at the middle of the interaural line (ear bar zero; EBZ). While the NMT v1.2 provided a single template configuration, the NMT v2 provides each template in two FOVs (standard and “full-head”) and at two spatial resolutions (standard and “low-res”) to make the templates suitable for a variety of tasks. Due to improved data type formatting, every version of the NMT v2 has a smaller file size than the NMT v2, including the “full-head” NMT v2 which has a much larger field of view.

**Table 1.** Comparison of versions of the NIMH Macaque Template (NMT) This table summarizes the differences between the NMT v2 templates and their predecessor, the NMT v1.2 (see section 2.2.4 for differences between v1.2 and its re-calculated form, NMT v1.3). The primary differences between the NMT v1.2 and NMT v2 are the new percentile-based averaging method and the reorientation of the NMT v2 to a stereotaxic orientation. Compared to the NMT v1.2, NMT v2 offers greater flexibility in the selection of template field of view (FOV), resolution, and atlases while having a smaller file size. The NMT v2 also has improved contrast between tissue classes, fewer artifacts, and a cleaner boundary between brain and CSF (refer to Figure 2 for a detailed comparison). The low-resolution (“low-res”; filenames contain “_05mm” to denote spatial resolution) variant of the NMT v2 may be useful for standard fMRI processing and for efficient sharing of data on platforms such as Neurovault. The expanded FOV (“full-head”; filenames contain “_fh”) variant of the NMT v2 may be useful for aligning datasets that have vastly different FOVs or positioning from the NMT v2. Abbreviations: FOV = field of view, AC-PC = anterior commissure-posterior commissure horizontal plane, WM = white matter, scGM = subcortical gray matter, GM = (cortical) gray matter, CSF = cerebrospinal fluid, BV = blood vasculature

|  | NMT v1.2 | NMT v2.0 |
| --- | --- | --- |
| **Symmetry** | Asymmetric | Asymmetric<br>Symmetric |
| **Orientation** | AC-PC | Stereotaxic (Horsley-Clarke) |
| **Origin** | Anterior Commissure | Ear bar zero (EBZ) |
| **Population averaging** | Mean | 20th - 80th percentile mean |
| **Intensity range (arbitrary units)** | 0 - 9.59 | 0 - 1024 |
| **Alignment script** | NMT_subject_align | @animal_warper (AFNI) |
| **FOV (mm)** | 63.00 x 86.75 x 61.25 | 64.00 x 78.00 x 50.00 ("standard")<br>100.00 x 100.00 x 75.00 ("full-head") |
| **Resolution (mm)** | 0.25 isotropic | 0.25 isotropic ("standard")<br>0.50 isotropic ("low-res") |
| **Matrix dimensions** | 253 x 347 x 245 | 256 x 312 x 200 ("standard")<br>128 x 156 x 100 ("low-res")<br>400 x 400 x 300 ("full-head") |
| **Template space name** | (None defined) | NMT2 |
| **Accompanying atlases** | D99 (Reveley et al., 2017) | CHARM<br>SARM (Hartig et al., this issue)<br>D99 (Reveley et al., 2017) |
| **Tissue segmentation** | 4-class (CSF, GM, WM, BV) | 5-class (CSF, GM, scGM, WM, BV) |
| **Data type** | Float (32 bit) | Short (16 bit int) |
| **File size** | 71 MB | 23 MB ("standard")<br>41 MB ("full-head")<br>6 MB ("low-res") |
| **Availability** | AFNI | AFNI |

Additionally, compared to the NMT v1.2, the NMT v2 datasets have several noticeable regions of improved clarity and contrast. Figure 2 provides a direct comparison of the symmetric NMT v2 and the NMT v1.2. The delineations among GM, WM, and CSF, as well as the discrimination between dura and GM, are clearer in the NMT v2. Hyperintensities in the frontal and temporal lobes of the NMT v1.2, driven by the merging of artifacts outside brain tissue with GM, have been markedly reduced in the NMT v2. We also observe less blurring of “brain” tissue with “non-brain” tissue such as blood vasculature and cranial nerves. These improvements allow for more precise identification of brain structures, which should improve alignment to the NMT v2. The improvements are especially beneficial for automated tools for template-based brain masking, segmentation, and surface extraction that can be thrown off by regions without clear delineation of tissue boundaries.

**Figure 2.**
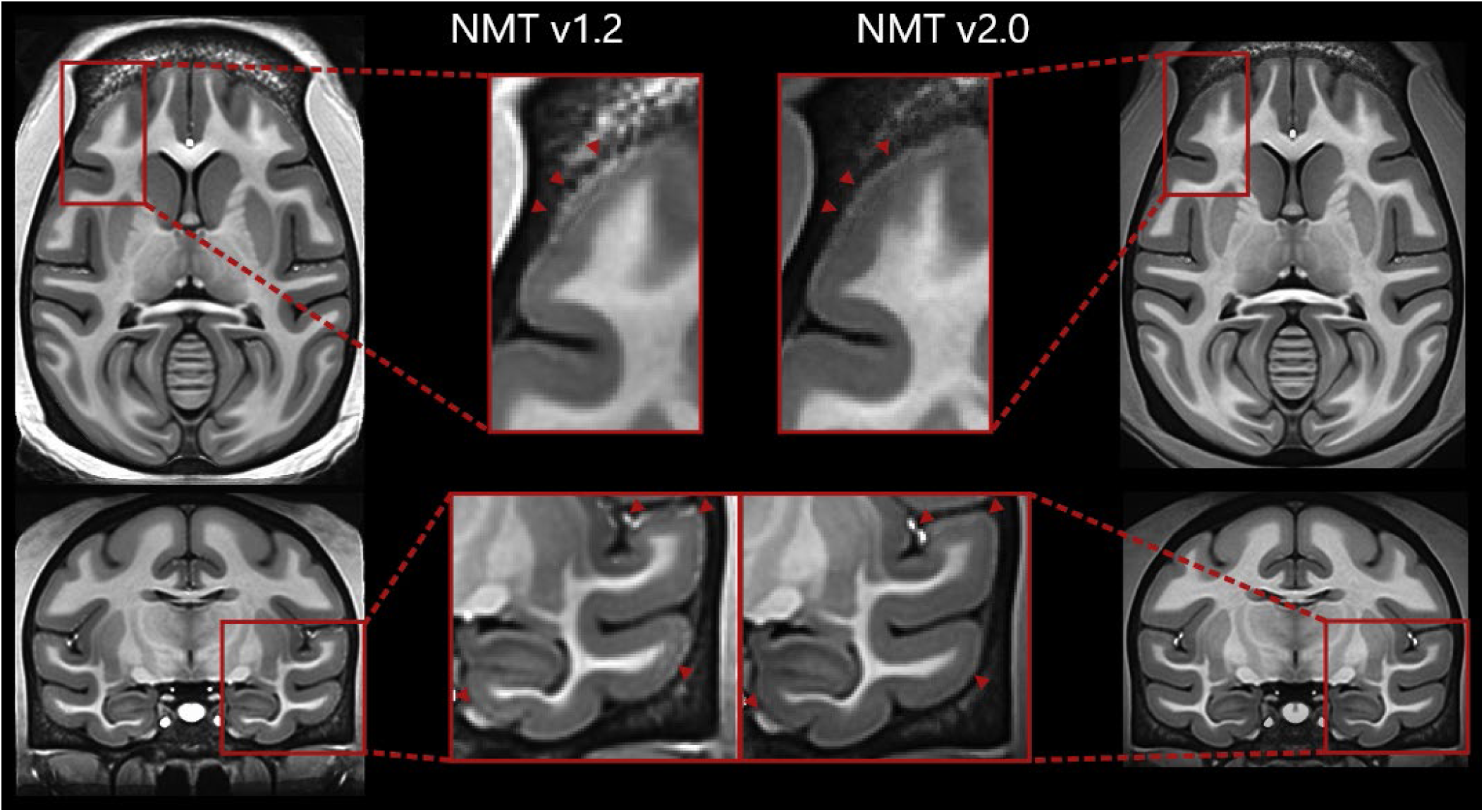
The NMT v2 has improved spatial contrast and structural delineation. The delineation of brain structures is directly compared in the NMT v1.2 (left) and the symmetric NMT v2 (right). Hyperintensities driven by sinus cavities previously resulted in blurred boundaries of the frontal surface of the NMT v1.2 (red box; top left). These hyperintensities in GM have been largely eliminated in the NMT v2 (red box; top right). In the temporal lobe (bottom), hyperintensities were also present in GM in the NMT v1.2 (red box; bottom left) and removed in the NMT v2 (red box; bottom right). We also observe improved delineation of “brain” tissues from “non-brain” tissues (such as blood vasculature and cranial nerves; bottom, on the ventromedial surface and dorsally in the lateral sulcus). Red arrows identify regions of improved delineation or removal of artifacts in the NMT v2. As displayed in the full coronal and axial images, we observe a more consistent intensity across the WM in the NMT v2. Note that in this figure the NMT v2 was rotated to match the AC-PC alignment of the NMT v1.2 to facilitate comparison between the two templates.

#### 3.1.2 Stereotaxic coordinates

Figure 3 demonstrates the stereotaxic orientation of the NMT v2 relative to the AC-PC aligned coordinate system of the NMT v1.2. The difference in the pitch angle between the AC-PC and stereotaxic orientations was determined by alignment of scans collected in a Horsley-Clarke stereotaxic apparatus to the NMT v1.2. Alignment required a mean pitch angle rotation of 12.3 deg, with a standard deviation of 1.5 deg (n = 10 monkeys). We therefore rotated the new template by this pitch angle, with rostral regions pivoting dorsally, to achieve the stereotaxic orientation of the NMT v2. The origin, which was at the center of the anterior commissure for NMT v1.2, has been moved for the NMT v2 to the intersection of the midsagittal plane and the interaural line. This is the origin of the Horsley-Clarke stereotaxic apparatus, where it is commonly referred to as ear bar zero (EBZ).

**Figure 3.**
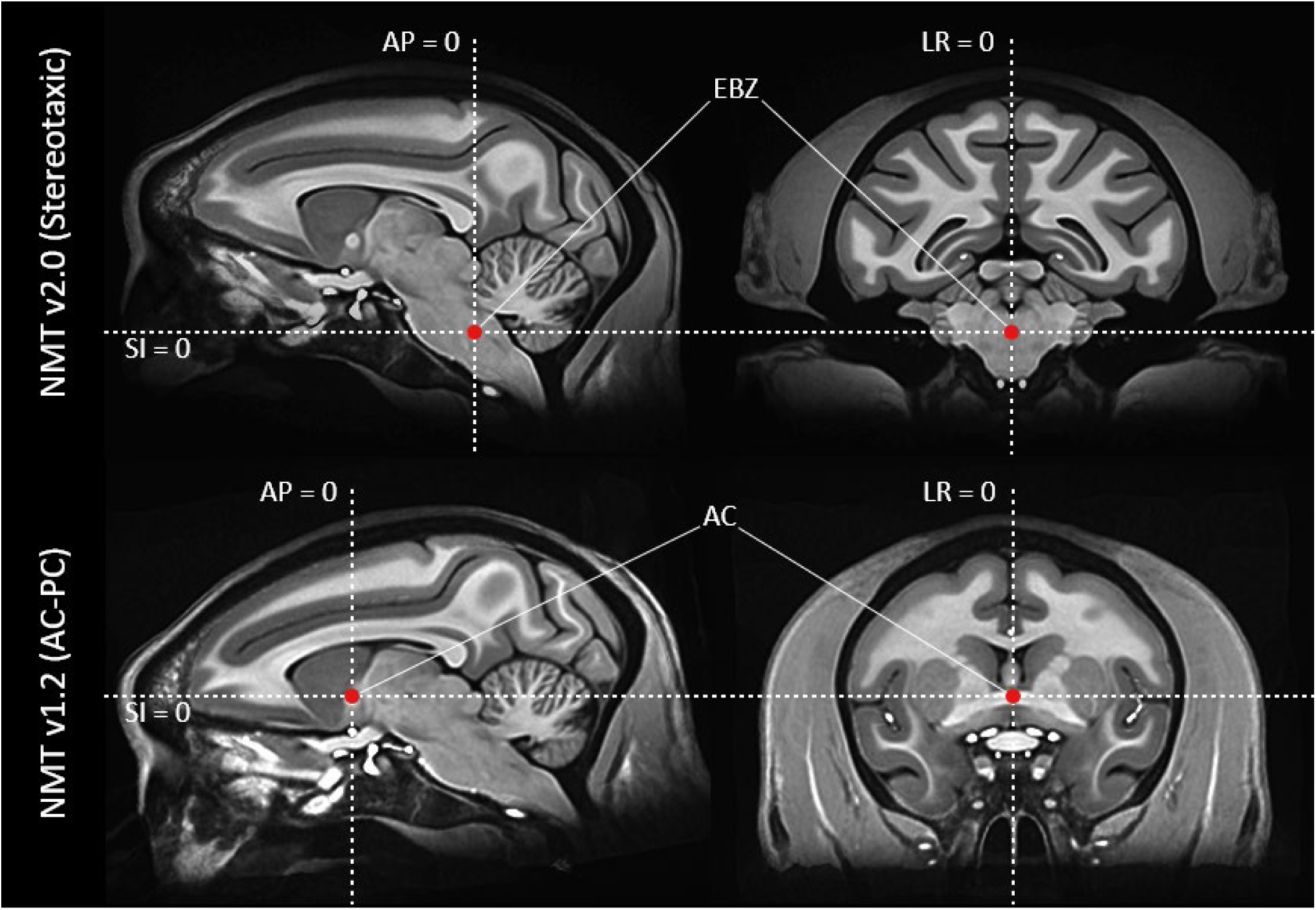
The orientation and coordinate system of the NMT v2. The orientation of the NMT v2 (top) is that of the Horsley-Clarke stereotaxic apparatus, which is commonly used to report coordinates in macaque research. The origin, denoted by a red dot, is at the midpoint of the interaural line, referred to as ear bar zero (EBZ). In contrast, the NMT v1.2 (bottom) is oriented so the horizontal plane passes through two internal brain landmarks, namely the anterior commissure (AC) and the posterior commissure (PC), or AC-PC alignment. The origin of the NMT v1.2 lies at the midpoint of the anterior commissure. Abbreviations: AP = Anterior - Posterior axis, SI = Superior-Inferior axis, LR = Left-Right axis.

The NMT v2 provides a useful guide for planning surgeries on a typical rhesus macaque. Preliminary surgical planning for a particular animal may be achieved by rigid alignment of the monkey’s pre-surgical MRI to the NMT v2, thus bringing the animal into approximate stereotaxic orientation while preserving the animal’s size and morphology. This can be achieved with @animal_warper by using the “-align_type rigid” (or “rigid_equiv”) option. However, the orientation of the animal’s head in the stereotaxic apparatus during surgery may differ slightly from this computed alignment. This deviation can be estimated from the inter-monkey variation in pitch angle (std = 1.5 deg) we observed in our own macaques. More precise determination of surgical coordinates can be achieved by collecting an MRI of the subject in a stereotaxic apparatus (preferably the one that will be used during surgery). However, this requires a deeper level of anesthesia and analgesia for the scan, as well as the availability of an MR-compatible stereotaxic device. Even then, scans collected from the same monkey in a stereotaxic apparatus on different days showed some intrasubject variation in the computed pitch angle (average std = 0.8 deg, n = 3 monkeys; 2 monkeys with 2 scan sessions and 1 monkey with 3 scan sessions). While tooth markers attached to the stereotaxic apparatus may be able to improve the reproducibility of head position in the device, this remaining variability suggests that there are inherent limits on the precision of presurgical planning.

#### 3.1.3 Masks and Segmentation

We provide detailed, manually refined masks with the NMT v2, including a brain mask and a cerebellum mask. Figure 4 shows the brain mask, 5-class tissue segmentation mask, and level 1 of the CHARM parcellation on the symmetric NMT v2 template. The same masks are displayed on the asymmetric NMT v2 in Figure S2. Researchers may apply these masks to data aligned with the NMT v2 templates for ROI-based analyses, tissue-based signal regression, masking, etc. Alternatively, the NMT v2 is useful for template-based segmentation and brain masking, as these masks may be transformed back to the undistorted anatomical scan (e.g., using @animal_warper), where they can be used to ascertain the native volumes of different structures and for ROI-based analyses.

**Figure 4.**
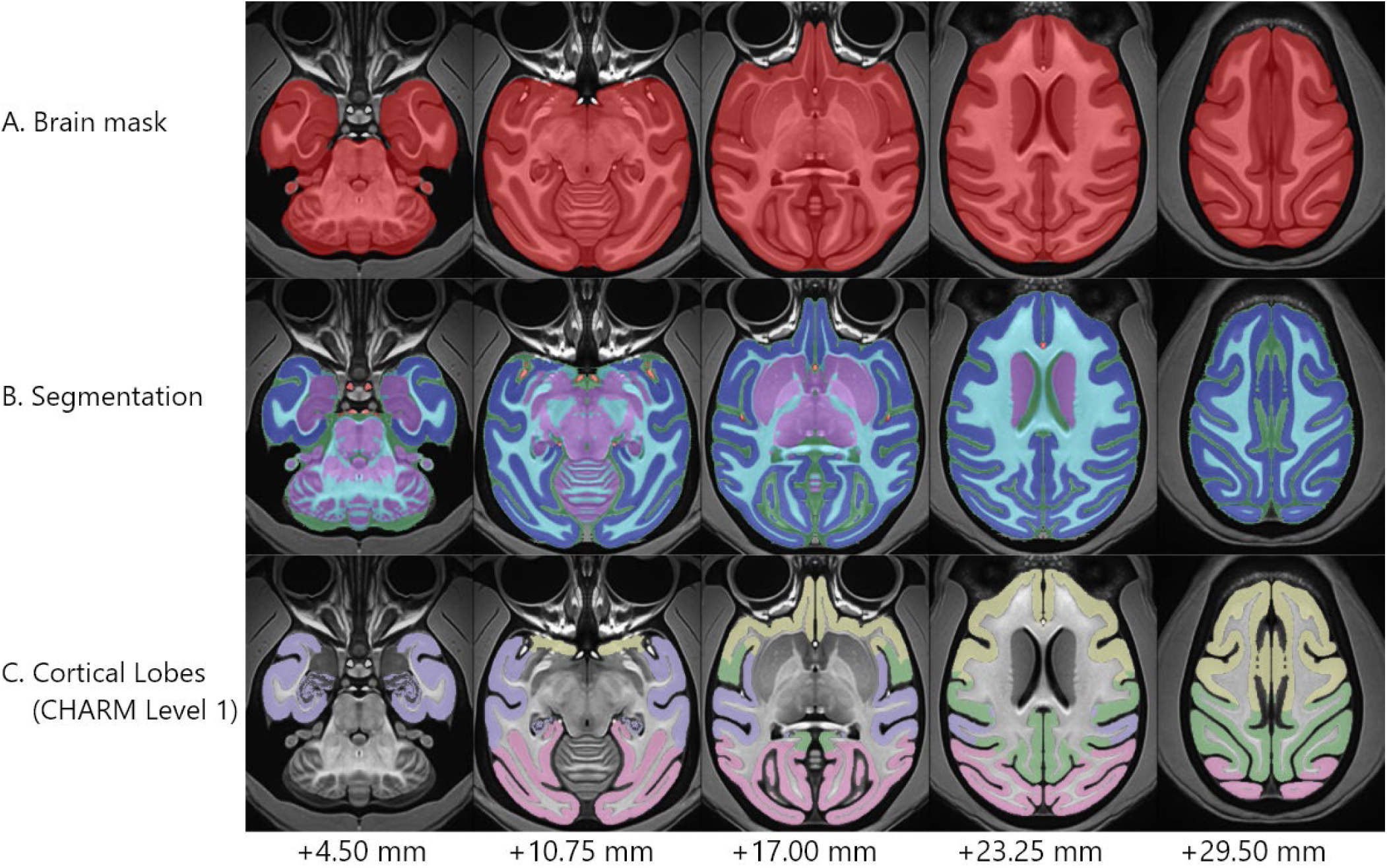
The symmetric NMT v2 and associated datasets. Axial slices through the symmetric NMT v2 are shown as an underlay, with various masks distributed with the template overlaid translucently. A) The symmetric NMT v2 brain mask (red) captures the brain and associated vasculature. B) The brain mask is further divided into a 5-class segmentation of the following types: cerebrospinal fluid (CSF; green), subcortical gray matter (scGM; purple), cortical gray matter (GM; dark blue), white matter (WM; light blue) and large blood vasculature (BV; red). C) Multiple atlases are provided with the NMT v2, including the CHARM. Level 1 of the CHARM hierarchy is displayed here, which consists of the frontal (yellow), temporal (purple), parietal (green) and occipital (pink) cortical lobes. Distances are superior to the interaural meatus. The asymmetric segmentation is shown in Figure S2.

#### 3.1.4 Size of the NMT

The volumes and mean intensity of the various tissue classes are provided in Table 2. We observe the volume of the brain mask to be 91.0 cc and the volume of the cerebellum to be 7.4 cc, which is consistent with our observations in the NMT v1.2 (Seidlitz, Sponheim et al. 2018) and other macaque brains (Lepage et al., this issue). The skull stripped symmetric NMT v2 template extends 29.375 mm laterally from the midline in both directions. In the anterior-posterior axis, the symmetric NMT v2 brain extends 47.000 mm anterior and 26.250 mm posterior to the interaural line. The NMT v2 runs from 39.000 mm superior to the interaural line to its cutoff point in the brain stem at 12.500 mm inferior to the interaural line. Both the anterior-posterior and medial-lateral extent agreed closely with the NMT v1.2 and the Paxinos atlas (Seidlitz, Sponheim et al. 2018; Paxinos et al., 2008). The superior-inferior extent of the NMT v2 was ∼5.5 mm greater than these other atlases because of its stereotaxic orientation and inclusion of more of the brainstem.

**Table 2.** Volumes of various tissue classes in the NMT v2 segmentation. The volumes of tissue classes and structures in the NMT v2 as determined from the NMT v2 masks. Template intensity values range from 0-1024 (arbitrary units). The “other” tissue class includes non-classified elements within the brain mask, including cranial nerves and dura. The cerebellum, which is part of the brain mask, is also shown separately. Abbreviations: A.U. = arbitrary units, CSF = cerebrospinal fluid, scGM = subcortical gray matter, GM = cortical gray matter, WM = white matter, BV = blood vasculature.

| Tissue Class | Volume (mm<sup>3</sup>) | Percentage | Mean (A.U.) | Intensity |
| --- | --- | --- | --- | --- |
| brain mask | 91,067 | - |  | 576 |
| CSF | 12,476 | 13.7 % |  | 301 |
| GM | 38,225 | 42.0 % |  | 537 |
| scGM | 15,732 | 17.3 % |  | 601 |
| WM | 23,405 | 25.7 % |  | 784 |
| BV | 158 | 0.2 % |  | 784 |
| other | 1,072 | 1.2 % |  | - |
| cerebellum | 7,371 | - |  | 620 |
| CSF | 99 | 1.3 % |  | 391 |
| scGM | 4,527 | 61.4 % |  | 555 |
| WM | 2,745 | 37.2 % |  | 737 |

#### 3.1.5 Surfaces

We provide high-resolution cortical surfaces of the NMT v2.0 templates using the new CIVET-macaque cortical surface reconstruction pipeline (Lepage et al., this issue). Surfaces include pial, WM (at the GM-WM boundary) and mid-cortical (between the pial and WM) surfaces, and are provided in the standard GIFTI format, allowing them to be used in most neuroimaging surfaces viewers, including AFNI’s SUMA visualization interface (Saad et al., 2004). These surfaces have per-vertex correspondence between white and pial surfaces, and the nodes are standardized between subjects (for more details about standard meshes, see Saad and Reynolds, 2012 and Lepage et al., this issue). These surfaces may be used to project statistical maps, ROIs, or other data for visualization. In Figure 5 and Figure 6, for example, we project multiple levels of the CHARM onto the mid-cortical surface, as displayed in SUMA. All available surfaces are displayed in Figure S3. Surfaces generated through CIVET-macaque may be saved in the NIFTI/GIFTI format and made compatible with SUMA using a new AFNI command, @SUMA_Make_Spec_CIVET. Note that while we provide these surfaces for visualization purposes, we recommend surface-based analyses be performed by processing your data through CIVET-macaque and projecting your functional data to a native surface that has been standardized to match the CIVET NMT average surface (“rsl” files in the CIVET output). The CIVET NMT average surface was generated through surface-based nonlinear registration of 29/31 of the NMT v2 volumes. For group analysis, data projected onto this average surface through CIVET-macaque may be averaged at the node level, avoiding blurring and topology concerns associated with volumetric data. 3D-printing-compatible files of all NMT v2.0 surfaces are included so that these surface objects may also be brought into the real world. Printed models of these surfaces may be useful for demonstration purposes and for surgical planning.

**Figure 5.**
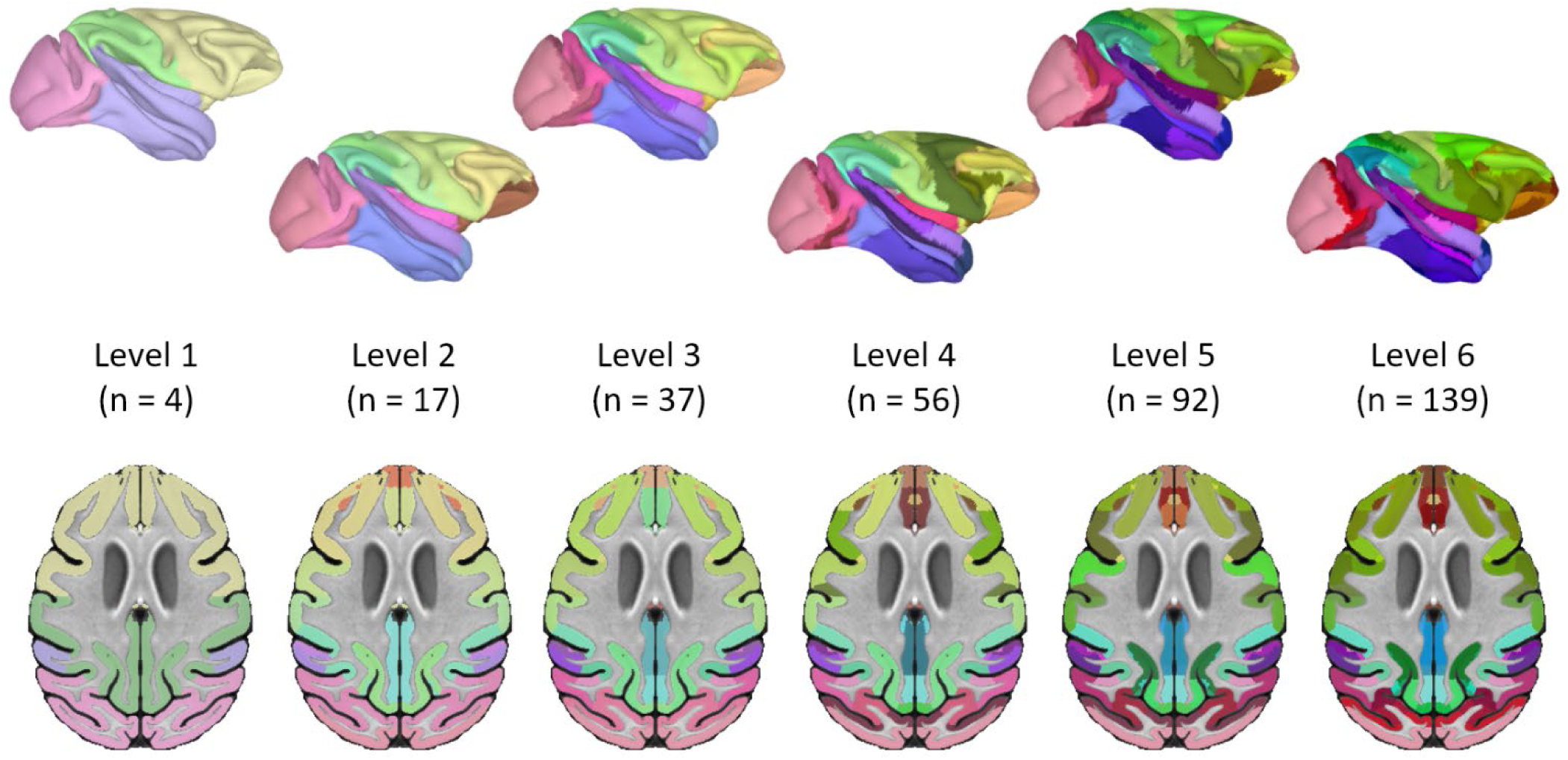
The Cortical Hierarchy Atlas of the Rhesus Macaque (CHARM). The CHARM parcellates the macaque cortical sheet at multiple spatial scales. Level 1 of the CHARM (left) contains the four cortical lobes. As the CHARM level increases, the cortical lobes are increasingly subdivided based on anatomical landmarks, functional relationships and cytoarchitectonics. The final level (level 6; right) consists of regions derived from the D99 macaque atlas, with exceptionally small D99 areas merged into neighboring regions to create ROIs robust to regridding of the atlas to individual scans. Each of the six levels of the CHARM are shown (along with the number “n” of regions per level) both projected onto the right hemisphere of the symmetric NMT v2 mid-cortical surface (top) and in an axial slice (24.5 mm superior to the origin) of the symmetric NMT v2 template (bottom). The color scale shows hierarchical relationships between levels, with each ROI sharing a similar hue to its related ROI in the level above. As the CHARM level increases, the saturation of the ROI color increases in kind. The mid-cortical surfaces were generated using CIVET-macaque (Lepage et al., this issue) and displayed using SUMA (Saad et al., 2004).

**Figure 6.**
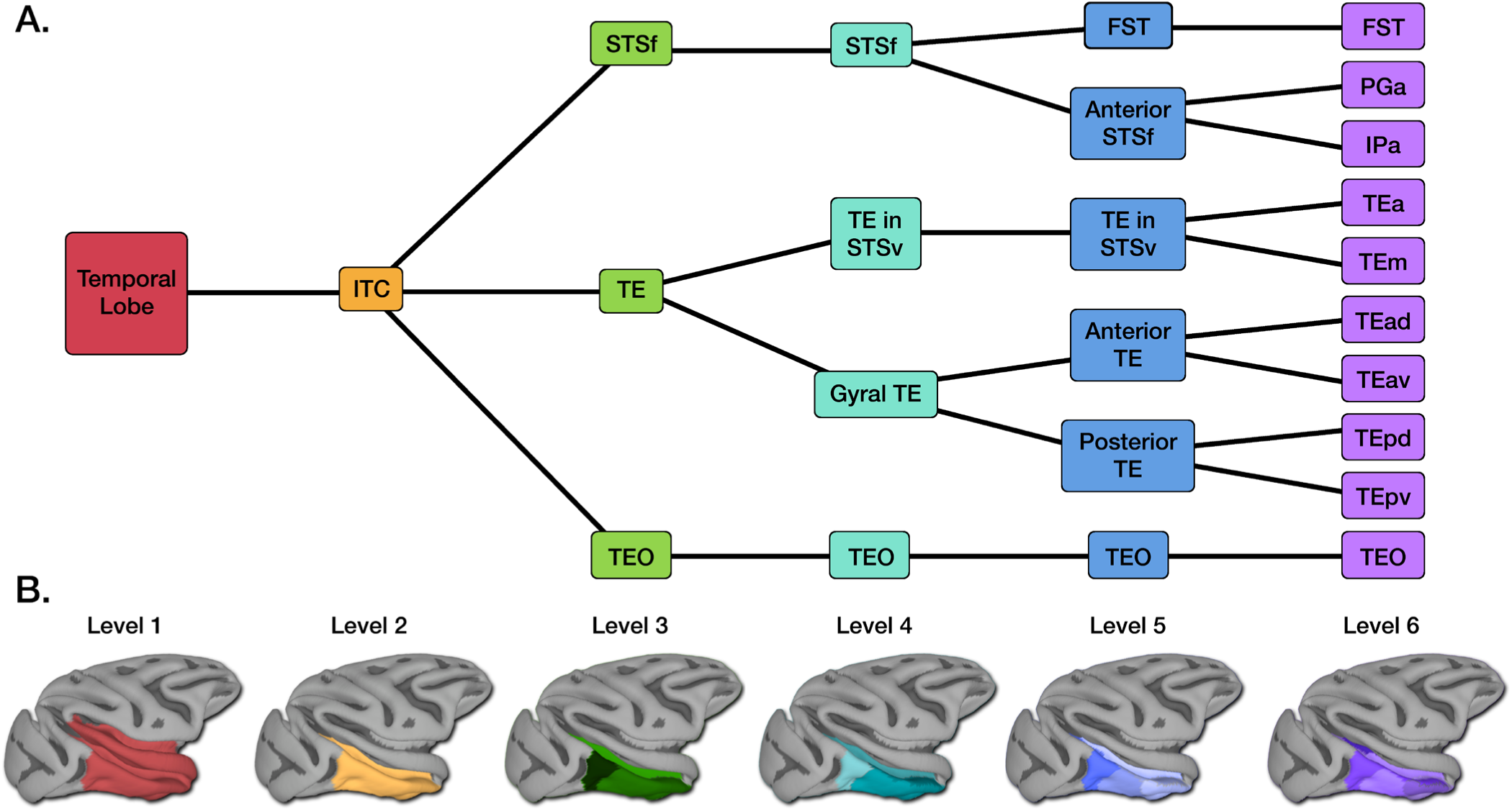
Example of the CHARM hierarchical structure. A) Tree diagram showing successively finer parcellation of the inferior temporal cortex (ITC). B) Lateral views of the symmetric NMT v2 surface with CHARM regions displayed in color. The ITC (Level 2, orange) is a visual region within the temporal lobe (Level 1, red). Other temporal lobe regions are not depicted. Levels 3-6 show the ITC subdivided into progressively smaller subdivisions. In level 3 (green), the ITC is subdivided into areas TE, TEO, and the fundus of the superior temporal sulcus (STSf). In level 4 (cyan), area TE is split into its portions on the ventral bank of the superior temporal sulcus (STSv) and its lateral and ventral portion on the middle and inferior temporal gyri (gyral TE). In level 5 (blue), both the gyral portion of area TE and the fundus of the STS are split along the anterior-posterior axis. Level 6 (purple) preserves the D99 parcellation. Area TE is comprised of 8 parts, with both anterior and posterior area TE further splitting into dorsal and ventral subdivisions and the STSv splitting into its cytoarchitectonic subdivisions (TEa and TEm). At this level, the STSf is divided into areas IPa and PGa and the floor of the superior temporal area (FST). The mid-cortical surfaces were generated using CIVET-macaque (Lepage et al., this issue) and displayed using SUMA (Saad et al., 2004).

### 3.2 The Cortical Hierarchy Atlas of the Rhesus Macaque (CHARM)

The CHARM describes the anatomical makeup of the cortical sheet at 6 different spatial scales (Figure 5). The broadest scale (CHARM level 1) contains the four cortical lobes. In the subsequent levels, the cortical lobes are iteratively subdivided until the finest scale (CHARM level 6), which has 139 regions. Across its 6 levels, CHARM has a total of 246 unique regions, with some regions appearing on multiple levels. This hierarchical organization allows researchers to use ROIs that best suit their data. For example, ROI-based analyses of fMRI data may require large ROIs that do not disappear when resampled to a coarse EPI resolution; lower levels of the CHARM are suitable for these analyses. The CHARM also allows researchers to select ROIs that best support their *a priori* hypotheses; while level 6 of the CHARM parcellates area TE into 8 subdivisions, someone investigating area TE as a whole may select all of area TE as a single ROI at level 3. Table 3 further describes the levels of CHARM, demonstrating how as the CHARM level increases, the number of ROIs increase, and the spatial scale of these ROIs decrease. Figure S4 demonstrates how individual voxel labels change with CHARM level for some example regions. A list of regions at each level of the hierarchy, and a table showing the transitions between regions at different CHARM levels is provided as a CSV file with the atlas file in the NMT v2 package.

**Table 3.** Basic characteristics of the CHARM hierarchy. The six levels of CHARM represent the organization of the macaque brain at different spatial scales. The lower levels of CHARM parcellate the cortex using a few, large regions of interest (ROIs), while higher levels organize the brain into smaller, more numerous ROIs; these higher levels still reflect the structure of the lowest ones, hence the atlas is hierarchical. The mean volume across ROIs in each level is listed as is the 25^th^ and 75^th^ percentile. Note that some ROIs may be present on multiple CHARM levels. The full table of CHARM values is provided as a CSV file in the NMT v2 distribution.

| Level | # of ROIs | ROI Vol., mean (mm<sup>3</sup>) | ROI Vol., 25-75% (mm<sup>3</sup>) |
| --- | --- | --- | --- |
| 1 | 4 | 9,717 | 8,722 - 10,469 |
| 2 | 17 | 2,286 | 1,205 - 3,332 |
| 3 | 37 | 1,051 | 521 - 1,188 |
| 4 | 56 | 694 | 261 - 897 |
| 5 | 92 | 422 | 125 - 543 |
| 6 | 139 | 280 | 76 - 302 |

#### 3.2.1 An example of the CHARM parcellation

Figure 6 shows an example of the CHARM parcellation of the inferior temporal cortex (ITC). The ITC (level 2) is a visual region involved in shape processing and object recognition that comprises much of the ventral temporal lobe (level 1). At level 3, ITC is divided into its main cytoarchitectonic and functional regions, areas TE and TEO, and a third region, defined geographically as the fundus of the superior temporal sulcus (STSf) (Bailey and von Bonin, 1947; Kravitz et al., 2013; Saleem and Logothetis, 2012). Area TE is partitioned at level 4 into the portion on the ventral bank of the STS (STSv) and the portion on the lateral and ventral surfaces (gyral TE). The former consists of two cytoarchitectonic subdivisions (TEa and TEm). Gyral TE splits into anterior and posterior TE (level 5), which are each further separated into dorsal and ventral subdivisions at level 6 (TEad, TEav, TEpd, TEpv) (Saleem et al., 2007; Saleem and Tanaka, 1996; Yukie et al., 1990). Area TEO is not further divided. At level 5, STSf is parsed into the more posterior floor of the superior temporal area (FST) and the anterior STSf, which breaks into areas PGa and IPa at level 6. In this way the ITC is progressively fractionated until at level 6 it has 10 constituents, all delineated by nonlinearly warping the D99 atlas (Reveley et al., 2017; Saleem and Logothetis, 2012) to the NMT v2. Additional examples of CHARM parcellation are shown in Figure S4.

### 3.3 Nonlinear warping with @animal_warper

At present, there are 6 total anatomical datasets in the AFNI macaque demos (each also available from PRIME-DE), spanning 3 different scanning centers and acquisition protocols (see demos for details). Each anatomical scan has varied FOV, brightness inhomogeneity, data quality, and angle of subject in the scanner. Nonlinear warping from each anatomical to the NMT v2 is estimated using @animal_warper, using default settings; the “low-res” versions of the NMT v2 are utilized in the demos by default to reduce runtime and file size (but the standard spatial resolution datasets are also distributed there, and the scripts can be adjusted by simply changing the input filenames). In all cases, the alignments to the standard template were judged to be successful with visual verification using the automatically generated QC images. Figure 7 shows axial images of the NMT v2 alignment to each subject’s input anatomical in native space (top row).

**Figure 7.**
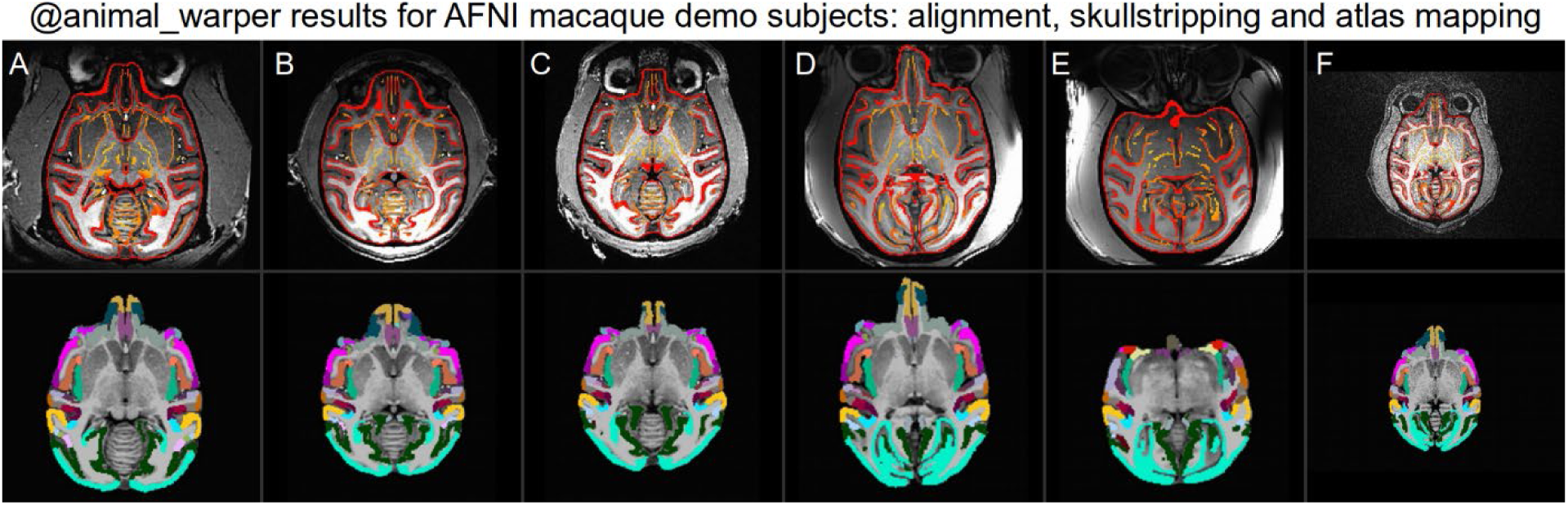
Results of alignment with @animal_warper for all subjects in the AFNI macaque demos. For each subject, the top row shows the edges of the NMT v2 template overlaid on the input anatomical, and the bottom row shows the warped CHARM level 5 ROIs overlaid on the skull stripped anatomical (after spatial normalization of brightness). Each image shows a middle axial slice, but each is in native space, so locations and scale vary due to size, shape, and angle of the subject’s head. In all cases @animal_warper’s alignment, skull stripping, and atlas-mapping appear to have been performed well. These anatomicals are from the MACAQUE_DEMO_REST (awake/anesthetized resting state fMRI) demo and available from PRIME-DE: subjects A-C are from NIMH (Messinger et al.); D-E are from NIN (Klink and Roelfsema); and F is from SBRI (Procyk, Wilson and Amiez); data from subject A is also used in the MACAQUE_DEMO_2.0 (task-based fMRI, with the same anatomical volume).

Several “follower” datasets in the NMT v2 template space are included in the @animal_warper command (e.g., the CHARM, D99 atlas, tissue segmentation and brain mask dataset), and the estimated nonlinear warp is applied to transform each of these into native subject space. Figure 7 (lower panel) shows an example of the placement of one follower set (CHARM, level 5) for each of the macaque demo subjects. The underlaid anatomicals have been skull stripped using the template’s brain mask warped to the native scan and intensity “unifizing” (bias correction) was applied to reduce the inhomogeneities resulting from scanner coil sensitivity. Figure 8 shows a greater selection of these follower datasets in native space (left columns), as well as surfaces generated from the warped atlas regions (right columns).

**Figure 8.**
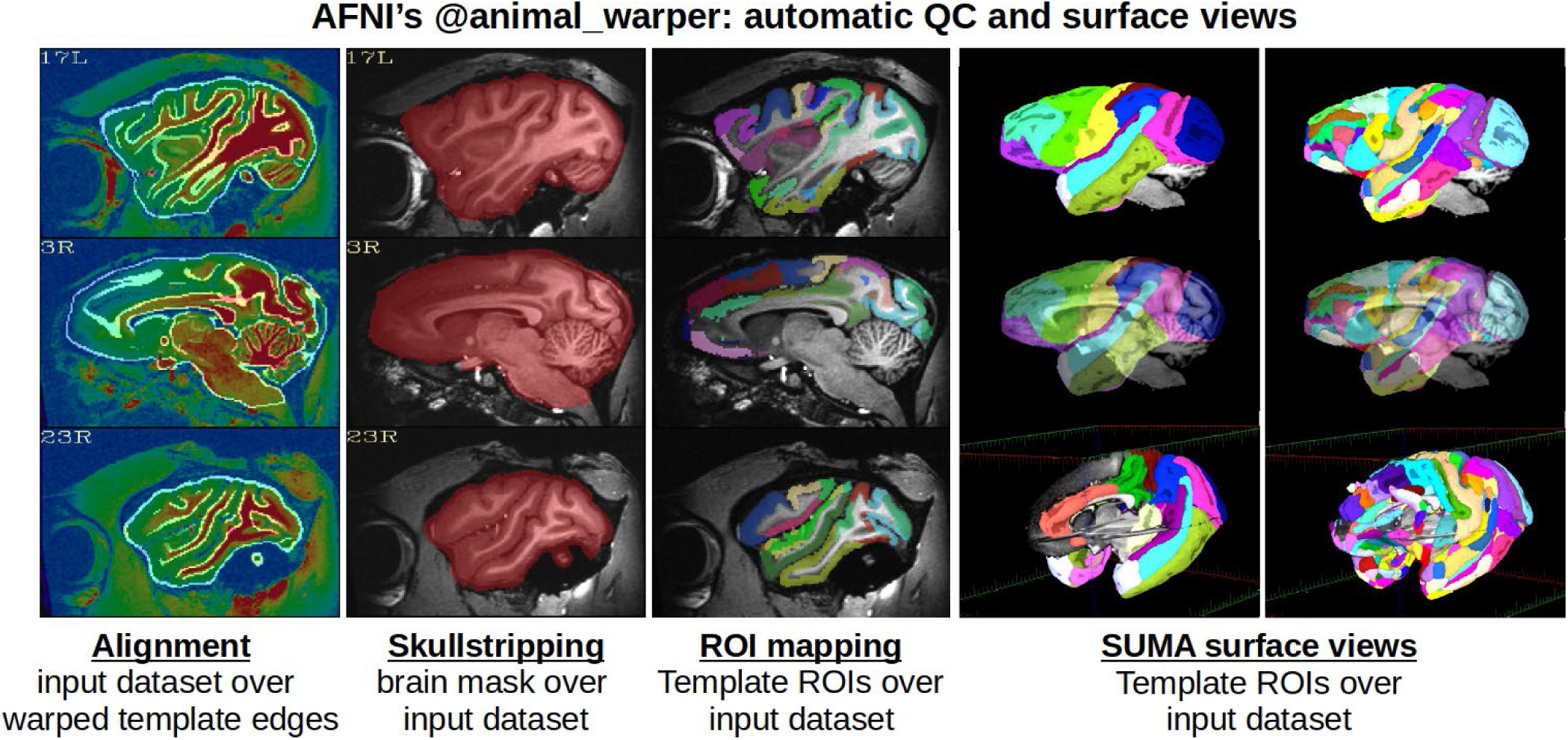
Examples of output datasets from @animal_warper. Outputs from @animal_warper for a single macaque subject in AFNI’s MACAQUE_DEMO_2.0 are shown. The first three columns show a subset of the automatic quality control (QC) output images created by the program, from left to right: alignment quality of the NMT v2 reference template (white edges of tissue boundaries underlay) to the original input dataset (translucent color overlay); skull stripping (warped brain mask overlaid on the input dataset); and atlas ROIs from the template space transformed to the input dataset (here, the CHARM, level 3). The fourth and fifth columns show surface views of the mapped CHARM Level 2 and Level 5 ROIs, respectively, with select slices of the subject’s anatomical scan using SUMA; these individual area surface files are automatically generated by @animal_warper, and can be viewed in different orientations, transparency levels and ROI combinations.

### 3.4 fMRI processing streams from afni_proc.py

#### 3.4.1 Task-based fMRI

The MACAQUE_DEMO_2.0 is a task-based fMRI study that uses data from a face patch localizer in a single subject; 13 total EPI runs were collected, and the demo scripts provide examples of analyzing either 4 or all 13 of the included EPI runs (with the former case for speed). Figure 9 shows a selection of the results from afni_proc.py’s QC HTML. The full F-stat of modeling (which can be interpreted as the variance of the model components divided by the variance of the residuals) in the top panel shows locations where the model is best fit to the data (degrees of freedom = 2, 1406). The given task was primarily visual (i.e. viewing images), so the predominance of high statistical values throughout the visual areas makes sense and would give the researcher confidence in their model setup, subject alertness and participation, and processing script design. Graphs of motion and outliers (Figure 9, middle panel) show that the subject had fairly small motion throughout (perhaps increasing slightly in the later runs) and correspondingly few volumes of high outlier fraction; this provides a complementary check to the F-stat (which includes all variance explained by the model, including both regressors of interest and nuisance regressors such as motion). Further assurance checks are included in the “warnings” section of the QC (Figure 9, lower panel), where it is noted that the motion peaks did not occur more frequently during a particular stimulus type (e.g. faces). This provides assurance that the effect size estimates (beta weights or coefficients) are not conflated with motion. The demo’s complete HTML output includes additional QC items, not shown here, including alignment quality, left-right consistency within the datasets, and more.

**Figure 9.**
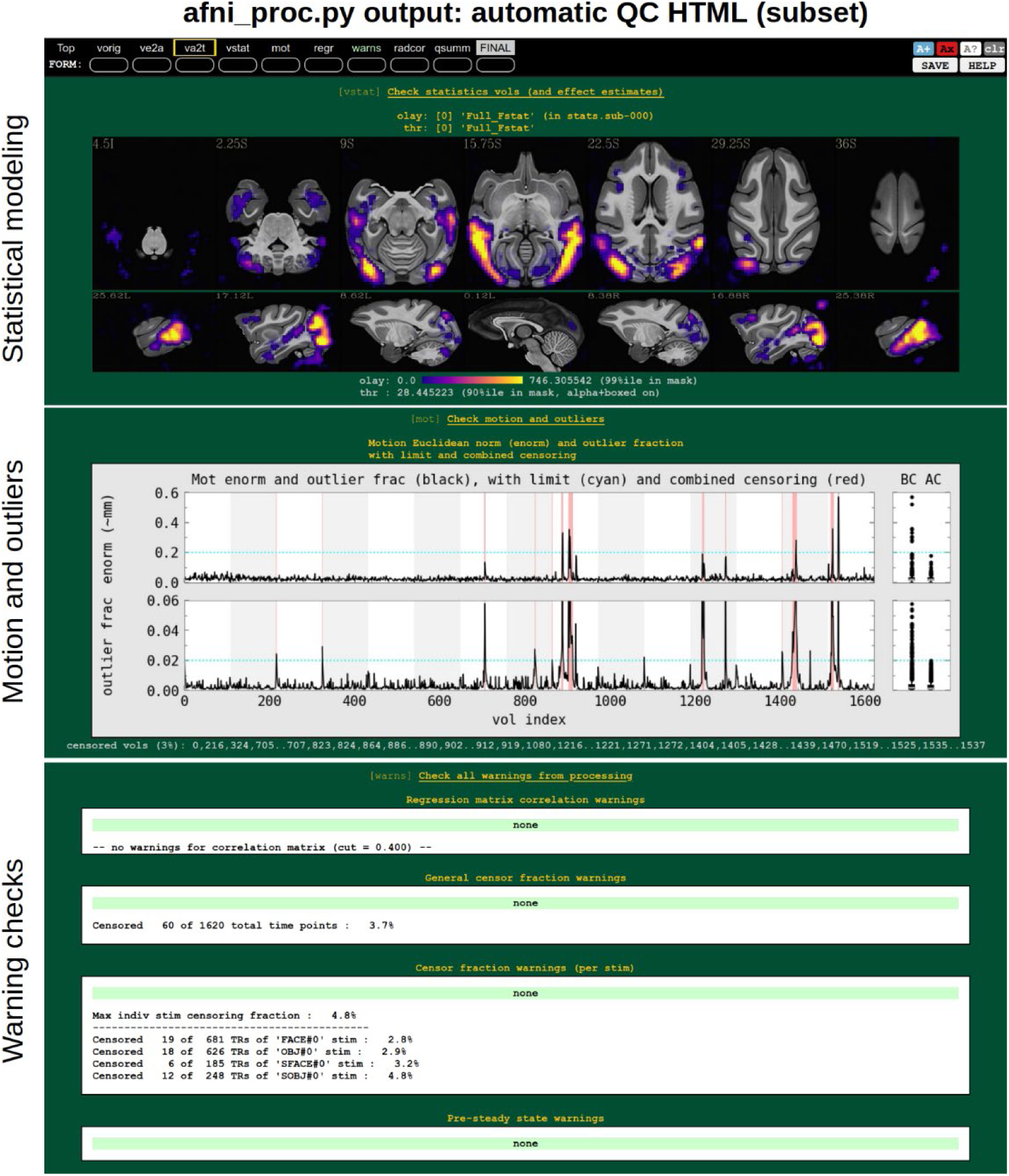
Some quality control (QC) steps provided by the afnI_proc.py fMRI analysis pipeline. Output is automatically generated in HTML format for efficient review in a browser. This example is from the task fMRI MACAQUE_DEMO_2.0 in AFNI. In the top block, the F-stat overlaid on the NMT v2 symmetric template shows brain regions where the modeled variance is much greater than residual variance, i.e., where the model specification is highest; in this case, visual areas involved in face, object, and scrambled stimulus perception. The next displayed block shows the motion (Euclidean norm of rigid body motion estimates) and the fraction of outlier voxels in the volume at each time point (x-axis), with censoring threshold levels for each (cyan line) and the resulting censored time points (red vertical line). The alternating gray/white background shows where EPI runs were concatenated (here, 15 runs), and the histograms at the right show the time-averaged parameter estimates before and after censoring (BC and AC, respectively). In the bottom block, warnings that afni_proc.py checks for while processing are presented, such as collinearity in the design matrix, total fraction of censored time points, fraction of each stimulus censored and pre-steady state volumes; warning levels are shown with a word and color above, and the level of greatest severity is also shown in the ‘warns’ tab in the HTML. Additional QC steps provided in the afni_proc.py output, but not shown here, include images of the raw data, alignments, local correlation structure and quantitative summaries.

Once the individual processing steps have been thoroughly checked for quality (i.e. the data does not appear to be corrupted by major artifact, misalignments, user errors such as wrong filenames, etc.), then one can proceed to view and interpret the results of interest. In the demo, 4 stimulus types are presented and there are various potential contrasts of interest that can be investigated. For example, the researcher might contrast an intact stimulus with its scrambled equivalent to identify brain regions that preferentially respond to an intact stimulus image over just the local features such as color and luminance that are also present in the scrambled image. As an example, Figure 10 shows the contrast of “intact faces vs scrambled faces.” For full results reporting, beta weights are shown (hot colors, intact>scrambled; cold colors, intact<scrambled), with thresholding applied to the statistical value (Chen et al., 2017). To better evaluate the modeling throughout the brain, display of the thresholded t-stat is not all or nothing: regions with |t|>3.30 (DF=1406, two-sided p<0.001; see Chen et al., 2019) are opaque and outlined in black, while sub-threshold regions are increasingly transparent. Additionally, to better evaluate the quality of modeling and guard against artifacts, the brain mask is not applied at this point in the analysis.

**Figure 10.**
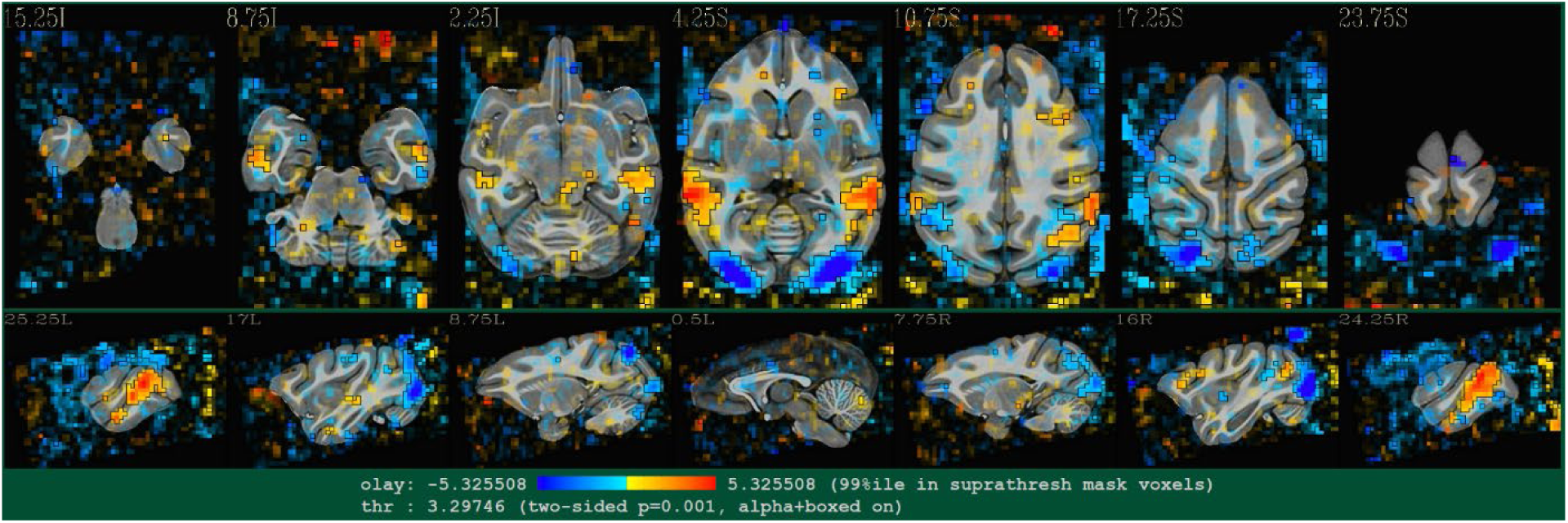
Example of a task-based fMRI contrast map generated using the afni_proc.py analysis pipeline. The contrast is of blocks where the monkey fixated centrally presented intact monkey faces versus scrambled monkey faces (i.e., “intact faces vs scrambled faces”). The overlay shows the effect estimates or beta weights (hot colors, intact>scrambled; cold colors, intact<scrambled) superimposed on axial and sagittal slices through the NMT v2 symmetric template. The t-statistic (DF=1406) is used for thresholding, applied in a way to show all relevant results from modeling: suprathreshold regions (two-sided p<0.001) are opaque and outlined in black, while values of lower significance are shown with increasing transparency. Because these data are from a single session of one subject, the brain region is not masked (to allow checking for artifacts, ghosting, motion effects, etc.).

Regions that respond significantly more to intact faces than scrambled faces are said to be face responsive. Regions that are both face responsive and face selective (i.e., respond significantly more to faces than other intact stimuli, such as objects) are referred to as face patches (Hadj-Bouziane et al., 2012, 2008; Tsao et al., 2008). Our task fMRI processing pipeline (Figure 10) identified face responsive regions bilaterally in the temporal lobe, which is the primary location of the macaque face patch system (Tsao et al., 2008). The early visual cortex, which responds well to oriented contrast edges, responded preferentially to the scrambled over the intact face stimuli. Note that the activation patterns shown in the demo are for a subset of the data collected during one scan session on a single subject to keep file sizes manageable. In practice, a typical analysis might involve many more repetitions within (or across) sessions, and possibly a multi-subject analysis.

#### 3.4.2 Resting state fMRI

The MACAQUE_DEMO_REST shows the analysis of resting state fMRI data collected from a variety of subjects with different protocols. Besides differences in scanner parameters, three of the macaques were anesthetized whereas three others were awake and had the contrast agent MION in their systems. We include a variety of data in the demo to reflect the various protocols currently in use and to assess the versatility of the afni_proc.py processing pipeline.

The processing of resting state fMRI contains many of the same QC considerations as task-based fMRI (e.g., checking for distortions, alignment, motion and outlier censoring), but because no stimuli are available to be regressors of interest, the main output of interest is the residuals themselves. The primary concern in analyzing these residuals is accounting for motion or other artifactual contributions (e.g., due to B0 inhomogeneity or scanner coils), while removing a minimal amount of neuronal signal. In general, this requires checking both estimated motion parameters, as well as the volumetric images and correlations patterns in the EPI directly.

Figure 11 shows a selection of the afni_proc.py-specified processing results from two of the demo subjects. To highlight the importance of proper QC, we present subjects who had notably different scan quality: one subject with low-to-medium motion (sub-01, top) and one with high motion (sub-02, bottom). Note that neither of these subjects had been anesthetized, and each had been injected with MION prior to scanning. Combined thresholding of the “enorm” (Euclidean norm of motion parameters) parameter and outlier fraction (fraction of voxels in a brain mask whose raw values are outliers) lead to censoring of 22% of time points for sub-01 and 54% for sub-02; the former produced a “medium” warning message in the HTML output, and the latter, a “severe” one.

**Figure 11.**
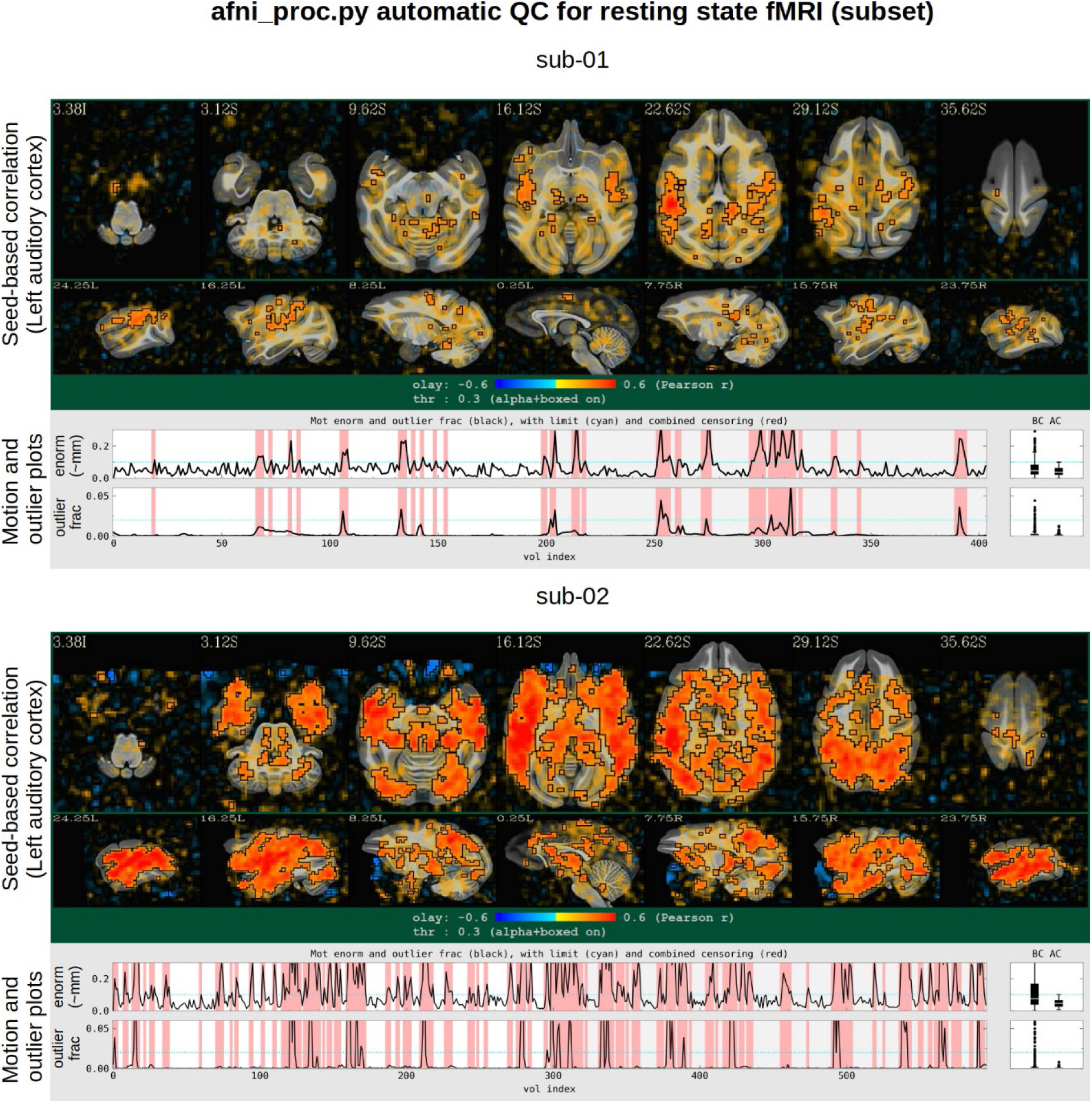
Resting state processing outputs for 2 of the subjects in the MACAQUE_DEMO_REST. For each subject, part of the automatic QC HTML created by afni_proc.py is shown: a seed-based correlation map (seed in the left auditory cortex at 20.1L, 5.0A, 21.0S in the symmetric NMT v2), with translucent thresholding and no brain masking applied; and the motion summary plots over the duration of the concatenated fMRI scans (“enorm,” the “Euclidean norm” of motion parameters; and “outlier frac,” the fraction of voxels in the brain mask that are outliers), with horizontal cyan lines showing censor limits and red regions showing censored time points. For sub-01, a fairly large fraction of time points have been censored (22%), producing a “medium” warning message in the HTML (not shown), but the seed correlation patterns have high left-right symmetry and have large values in physiologically reasonable locations. For sub-02, a very large fraction of time points has been censored (54%), producing a “severe” warning message (not shown), and indeed the correlation patterns appear to still be heavily influenced by motion (extremely high correlation throughout the brain, including non-GM tissues).

The *NMT2* space name is recognized within the QC-generating libraries of AFNI, and three seed-based (Pearson) correlation maps are generated automatically from locations spread throughout the macaque brain: the left auditory cortex, right primary visual and left posterior cingulate cortex (PCC). These seed locations have been chosen to provide information on major networks that show well-known activation patterns and that also span much of the cortex. Figure 11 displays the correlation map from the auditory seed, using translucent thresholding and no brain masking, as described in section 3.4.1 (for processing which included spatial smoothing). As might be expected from the motion plots, even after censoring, sub-02’s fMRI time series appear to be heavily affected by motion: correlations are unphysiologically high throughout the entire cortex, including in regions such as WM that are not expected to correlate with cortical GM. One also notices that the EPI FOV did not fully cover the whole brain, lacking coverage in the frontal region. Conversely, the pattern of correlation for sub-01 appears reasonable: fairly left-right symmetric, and with reasonably high values in expected areas, given the seed location. As a result, one would likely feel confident including sub-01 in further analysis, while excluding sub-02.

The single subject processing outputs from afni_proc.py can be combined in many ways for group analysis. One example provided in the demo is to make use of the CHARM for ROI-based analysis to investigate networks of interest. Figure 12 shows correlation matrices calculated with AFNI’s 3dNetCorr for sub-01 in levels 2-5 of the CHARM (with time series averaged within each ROI but no additional spatial smoothing). Consider the occipital visual network in the lower right corner of the matrix, visible with increasing structure through the CHARM levels, as well as the somatomotor regions (motor and SI/SII in level 2; M1/PM, SMA/preSMA, SI and SII in level 3; etc.) and the lateral temporal lobe network (ITC and STG/STSd in level 2; TEO, TE, STSf, STGr/STSd and STGc in level 3; etc.).

**Figure 12.**
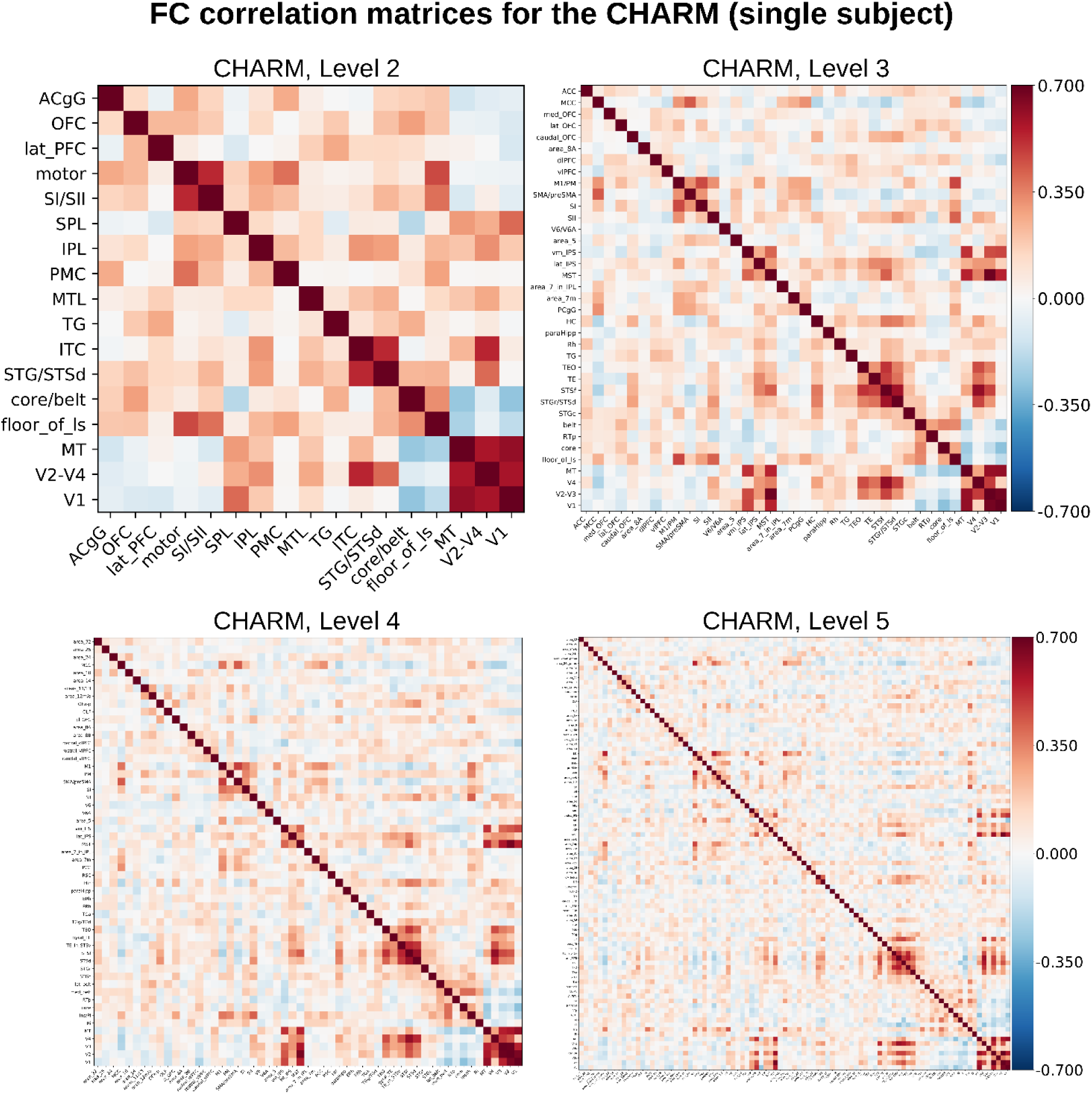
Resting state correlation matrices generated from a single subject using the CHARM. After the processing and regression modeling specified with afni_proc.py, one can combine the main output (i.e., the residual time series, in this case without spatial blurring) with an atlas (here, the CHARM) to conduct ROI-based analyses. We display the middle four CHARM levels as a demonstration of how spatial scale impacts resting state correlations. The hierarchical nature of the CHARM allows correlations to be evaluated at multiple spatial scales, allowing one to “zoom in” on details. For example, notice the clusters of correlation both within the visual regions (areas V1,V2-V4, and MT) and the somatomotor regions (motor, SI-SII), which are observable throughout all of the CHARM levels at various degrees of detail.

## 4. Discussion

We have described a new high-resolution anatomical template, the NMT v2, and a hierarchical cortical atlas (CHARM) that facilitate reproducible fMRI analysis in the macaque. These resources have been integrated into a comprehensive, start-to-finish processing pipeline for both task-based and resting state macaque fMRI data. The pipeline utilizes the AFNI commands @animal_warper to align fMRI data to a common space, and macaque-specific parameters in afni_proc.py to perform preprocessing and regression with quality control images.

### 4.1 The NIMH Macaque Template v2

The NMT v2 is a high-resolution template space for macaque neuroimaging, complete with masks, segmentation, and atlases. Its predecessor, the NMT v1.2, compared favorably to Rhesus macaque templates available at that time (Seidlitz, Sponheim et al., 2018) and no comparable template has been released since. Compared to this predecessor, the NMT v2 templates have superior contrast and definition, with fewer artifacts.

The NMT v2 provides a high-resolution target for alignment of macaque MRI data to a standardized space. Alignment is simple and accurate due to the fine anatomical detail of the NMT v2 and its integration with @animal_warper. Additionally, tissue masks, segmentation, and atlases can be used to process scans and assist in ROI-based analyses. For example, WM and CSF masks can be used in resting state analyses to control for non-neuronal sources of correlated activity. Similarly, CHARM can be used for ROI-based functional connectivity analysis (see section 4.2). For processing data in native space, these resources may be warped back to the original anatomical scans, where they can be used for template-based skull stripping, segmentation, and ROI-based analysis.

Presenting and reporting results in the NMT v2 space facilitates reproducible science and comparisons across time, subjects, and laboratories. Standardized coordinates obtained from NMT v2-aligned datasets enable the assessment of consistency across monkeys, the generation of probabilistic maps, and the performance of meta-analyses. These benefits have been realized in human neuroimaging for years, where there is broad adoption of template spaces such as the MNI space (Fonov et al., 2009), and the presentation of statistical maps in common spaces in repositories such as Neurovault (Gorgolewski et al., 2015). We selected the Horsley-Clarke stereotaxic coordinate system for the NMT v2 templates to facilitate comparison between noninvasive techniques, like functional neuroimaging, and invasive techniques such as lesions, chemogenetics and electrophysiology. Researchers can use the stereotaxic template and coordinate system to align scans of their subjects for early-surgical planning, to leverage fMRI activation maps or the atlases packaged with the NMT to target brain areas for invasive study, and to report the coordinates of surgical and experimental procedures.

The symmetric form of the NMT v2 is useful for interhemispheric comparisons of MRI data. While much less pronounced than for the human brain, there are cortical asymmetries in the macaque population (Lepage et al., this issue; Xia et al., 2020), and macaque brain morphometrics remain understudied. When assessing morphometrics, data should be warped to a symmetric template, as this provides a non-biased target for such analysis (Watkins et al., 2001). Warping to an asymmetric template has the potential to confound individual asymmetries with those present at the population-level. Beyond making interhemispheric comparisons easier and more reliable, symmetric templates facilitate functional and structural analyses that simply are not possible with an asymmetric template (or in the native space of the subject, for that matter). Voxel-mirrored homotopic connectivity, for example, maps resting state connectivity at identical locations across hemispheres, and relies on a symmetric template to calculate homotopic correlations between corresponding voxels in the two hemispheres (Zuo et al., 2010).

At a practical level, symmetric templates can make the development of atlases, maps, and other tools for MRI research more efficient. For example, cytoarchitectonic atlases are often developed on a single hemisphere and mirrored onto the other hemisphere (Reveley et al., 2017; Saleem and Logothetis, 2012). This is the case for the new Subcortical Atlas of the Rhesus Macaque (SARM), which was refined on a single hemisphere of the symmetric NMT v2 and mirrored to the other, identical hemisphere (Hartig et al., this issue).

### 4.2 The Cortical Hierarchy Atlas of the Rhesus Macaque (CHARM)

We have described and demonstrated the utility of the Cortical Hierarchy Atlas of the Rhesus Macaque (CHARM), a novel six-level anatomical parcellation of the macaque cerebral cortex, where the cortical sheet is subdivided into finer and finer parcellations at each successive level. By subdividing the cortical sheet at multiple spatial scales, the CHARM provides flexibility and precision to researchers conducting ROI-based analyses. The atlas allows large cortical activations to be efficiently described in terms of the composite regions that span that territory.

Several atlases have been created for the macaque brain and mapped to an MRI, including the D99 atlas (Reveley et al. 2017; Saleem and Logothetis 2012) and the Paxinos Atlas (Paxinos et al., 2008). The CHARM leverages the fine cytoarchitectonic delineations of macaque brain tissue from the D99 atlas to create a cortical hierarchy, expanding the utility of these regions. The D99 atlas was defined on an MRI collected *ex vivo*. We have observed that differences in sulcal positioning make alignment between *in vivo* and *ex vivo* brains difficult, with regions frequently extending across sulcal banks. The CHARM, which is tailored to the *in vivo* NMT v2, presents an easier, more accurate alignment target for *in vivo* neuroimaging. The *ex vivo* Calabrese DTI atlas grouped Paxinos atlas areas into a 5-level hierarchy (Calabrese et al., 2015; Supplementary Table 2), with 9 composite regions spanning the entire cerebral cortex. The CHARM is defined as a volumetric dataset of 6 ROI maps in the NMT2 space, which can be used as an anatomical reference atlas or for ROI-based analyses, and is accompanied by a descriptive table, as well. We note that subcortical regions are absent from the CHARM. A hierarchical atlas of the subcortex, called the Subcortical Atlas of the Rhesus Macaque, was recently developed in an international collaboration that includes our group (Hartig et al., this issue). Like the CHARM, the SARM parcellates the brain at 6 spatial levels, and is provided with the NMT v2. We encourage researchers to combine the CHARM and the SARM for questions relating to cortical-subcortical interactions.

The diverse spatial scale of the CHARM allows for ROI-based analyses to be performed on data collected at a wide range of resolutions, including fMRI voxel resolutions. While the default NMT v2 has a 0.25 mm isotropic resolution, anatomical data is typically collected with voxels that are twice as big on a side (0.50 mm) and EPI data is typically collected with voxels that are 6 times larger (1.5 mm). Hence, a single anatomical voxel can house 8 (=2^3^) NMT v2 voxels and a single EPI voxel can house 216 (=6^3^) NMT v2 voxels. Resampling atlases onto these coarser grids can impact the size and structure of regions, sometimes resulting in the loss of small or thin ROIs or the formation of discontinuities. The finest level of the CHARM was defined to map cytoarchitectonic features onto macaque MRIs, and as such may not be appropriate for ROI-based averaging of large functional voxels. The CHARM provides a solution for ROI-based functional analysis by representing the entire cortical sheet at multiple spatial scales. Larger ROIs from the broader levels of CHARM may be more robust to resampling by reducing the risk of disappearing or becoming discontinuous.

CHARM regions facilitate the description of significant functional activations. Such activations are often reported using center of mass coordinates, but this fails to fully represent the data and is insufficient for replicating findings. The center of mass leaves out critical information about the shape and distribution of an activation. Furthermore, depending on the shape of the activation, the center of mass may not even be located within the significant cluster. Sharing of statistical maps alleviates these concerns, but this is not always feasible. Instead of reporting the center of mass coordinates, researchers can find the CHARM region that best represents their data, and report that region and the overlap with the cluster. Because the CHARM represents the cortex at multiple spatial scales, it is more likely a CHARM region (or regions) can concisely represent an activation map as compared to a non-hierarchical atlas.

Atlases can also facilitate various forms of network and group analysis. Most standard, voxel-wise analyses in neuroimaging are performed in a “massively univariate” fashion and hence require various corrective procedures for multiple comparisons (e.g., clustering), typically introducing a need for multiple semi-arbitrary thresholding parameters. Atlases can reduce the dimensionality of datasets by orders of magnitude, from tens of thousands (voxels) to hundreds or less (ROIs), thereby reducing the required statistical corrections. The different spatial scales of the CHARM allow researchers further flexibility to select a CHARM level *a priori* that limits the number of multiple comparisons. Additionally, recent statistical work in AFNI has focused on including all elements of a group analysis (e.g., voxels or ROIs) in a single model, so that corrections for multiple comparison are not required. These Bayesian methods are presently available in AFNI (e.g., programs *RBA* and *MBA*) for ROI-based analyses. They offer additional advantages, including containing built-in model validation, allowing for full results reporting, and not requiring arbitrary thresholding (Chen et al., 2020, 2019a). Having a hierarchical atlas such as the CHARM allows a great deal of flexibility for the researcher to perform such analyses on an appropriate scale for their study and experimental design. Regions may even be combined across levels. For example, the anatomical seed used for a resting state analysis can be a broad composite region, assuming all the areas within it are themselves highly correlated. The average time course over this composite region may then be correlated with finer regions outside it. This results in a cleaner average time course, and fewer functional connectivity correlations to calculate.

### 4.3 Macaque fMRI processing in AFNI

AFNI and the NMT v2 represent a complete framework for macaque fMRI analysis (from raw data checking through preprocessing, single subject regression, and group level statistics), particularly with the inclusion of the CHARM for ROI-based analyses and spatial reference. AFNI is designed around the idea of allowing researchers to design and carry out their specific analyses of interest, and afni_proc.py in particular can be used to create highly customized pipelines to fit a wide variety of experimental designs across task-based, resting state and naturalistic paradigms. The command has (relatively) succinct syntax so that it can be shared and/or adapted easily; the commented processing script and intermediate datasets are preserved for validation and QC purposes; and several automatic QC features contained within the processing were also noted. We have also presented the @animal_warper program for nonlinear alignment and mapping of ROIs between spaces, which contains additional features of its own, such as automatic QC image generation, scripts for viewing the created surfaces and data tables of ROI properties (such as both absolute and relative sizes) before and after warping. The @animal_warper program has shown itself to be effective at transforming structural datasets to the NMT template. We have used it to successfully align PRIME-DE datasets from several sites, including those used in this study, as well as other sites used in a previous study that included macaque data (Glen et al., 2020). In addition, the AFNI software package includes over 600 programs of basic imaging/signal processing, statistics, and data visualization (both in volumes and on surfaces). The NIFTI/GIFTI file formats allow for cross-software integration; several separate analysis and pipeline tools that are focused on macaque fMRI processing exist, both for general analysis and specific operations.

The PRIMatE Resource Exchange (PRIME-RE; Messinger et al., this issue) was created to provide a single resource that organizes and describes the growing number of NHP-specific imaging resources available. Information on our AFNI macaque pipelines may be found at the PRIME-RE website^16^. We hope that this resource exchange website will facilitate the sharing of ideas on analysis, benefitting the research community as a whole. The PRIME-RE also provides a valuable resource for creating new tools and methods. The AFNI codebase has grown a great deal over its history due to suggestions from users at conferences, on message boards, and in person. We continually look to add more functionality to the software. Thus, it is likely that macaque researchers will find existing tools in AFNI that meet their research needs and that our continuing discussions with researchers will foster the genesis of new software and analysis features.

AFNI has many additional applications that can be relevant for macaque neuroimaging beyond fMRI processing. For example, there are a large array of group level statistical programs for voxel-wise analysis and clustering, as well as ROI-based methods; see Chen et al., (2019a, 2013) and Cox (2019). Some of the software display capabilities with SUMA have been noted above, but there are many other useful SUMA-based features, including the ability to project volumetric data onto a surface during fMRI preprocessing within afni_proc.py; this feature integrates usefully with CIVET-macaque generated surfaces (Lepage et al., this issue). Additionally, there are tools for processing diffusion weighted imaging (DWI) data, particularly the FATCAT subset of AFNI tools (Taylor and Saad, 2013), which also integrate with the freely available TORTOISE software (Pierpaoli, et al., 2010). In particular, FATCAT was designed to combine fMRI and diffusion-based studies through network analyses (e.g., Taylor et al., 2016), and also contains tools relevant for functional studies.

### 4.4 Conclusion

We have presented a comprehensive macaque fMRI analysis pipeline in AFNI, with a new hierarchical atlas (CHARM) in the improved NIMH Macaque Template (NMT) v2 space. The NMT v2 and the CHARM, along with demonstrations of their use with AFNI’s functional processing pipelines, are currently available via the AFNI website^17^. Collaborative initiatives in NHP neuroimaging have progressed rapidly in the past few years, presenting the need for standardized coordinates spaces to share reproducible results, robust alignment programs for alignment of data to these standardized spaces, and customizable pipelines to conduct macaque fMRI preprocessing and analysis. As demonstrated through our pipelines, the NMT v2 and AFNI provide the macaque community with all these capabilities.

## CRediT authorship contribution statement

**Benjamin Jung:** Conceptualization, Methodology, Software, Formal analysis, Validation, Data Curation, Writing - Original Draft, Writing - Review & Editing, Visualization

**Paul A. Taylor:** Conceptualization, Methodology, Software, Formal analysis, Validation, Data Curation, Writing - Original Draft, Writing - Review & Editing, Visualization

**Jakob Seidlitz:** Conceptualization, Methodology, Writing - Review & Editing, Visualization

**Caleb Sponheim:** Conceptualization, Methodology, Writing - Review & Editing, Visualization

**Pierce Perkins:** Methodology, Writing - Review & Editing

**Leslie Ungerleider:** Resources, Writing - Review & Editing, Funding acquisition

**Daniel Glen:** Conceptualization, Methodology, Software, Resources, Writing - Review & Editing, Supervision

**Adam Messinger:** Conceptualization, Methodology, Resources, Validation, Writing - Original Draft, Writing - Review & Editing, Supervision

## Acknowledgements

The authors would like to thank Claude Lepage for generating cortical surfaces of the NMT v2 and for consultation on template creation, Rick Reynolds for his help in putting together the afni_proc.py commands for the macaque demos, and Richard Saunders for feedback on the CHARM.

We also thank David Yu for his help in the scanning procedures and data collection, and researchers in the Laboratory of Brain and Cognition and the Laboratory of Neuropsychology who contributed anatomical scans of their subjects to this project.

This work was funded in part by the Intramural Research Program of the NIMH and NINDS (ZICMH002899 and ZICMH002888). This work utilized the computational resources of the NIH HPC Biowulf cluster^18^.

## Supplementary Materials

**Figure S1.**
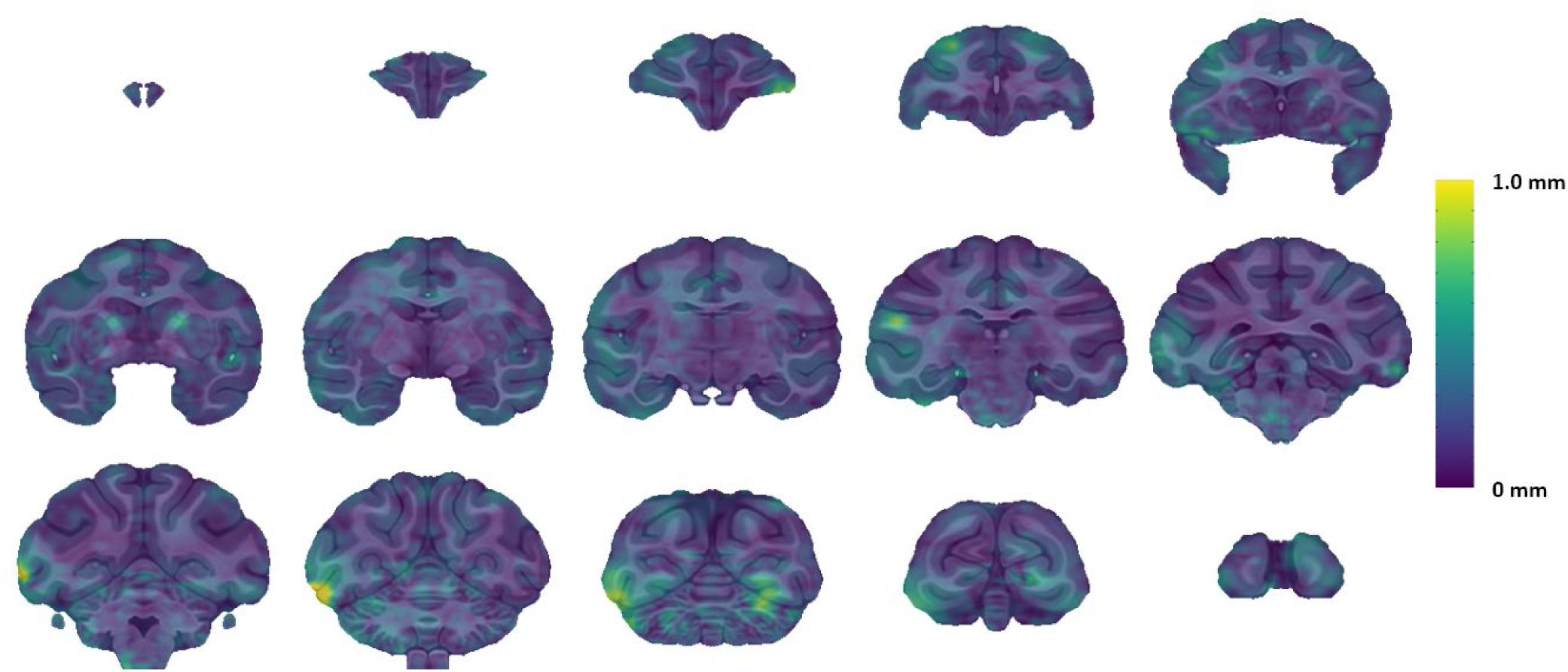
Magnitude of displacement (MOD) between the symmetric and asymmetric variants of the NMT v2. Morphological differences between the two variants of the NMT v2 are typically small (sub voxel, <0.25 mm). The MOD is the physical distance between aligned points in the two spaces, calculated as the L2 norm (i.e., the square root of the sum of squares) of the nonlinear warp field components at each point. The MOD median, stdev and maximum values are 0.18 mm, 0.12 mm, and 1.25 mm, respectively. Only 2.36% of voxels have MOD > 0.50 mm (two voxels) and only 0.04% of voxels have a MOD > 1.0 mm.

**Figure S2.**
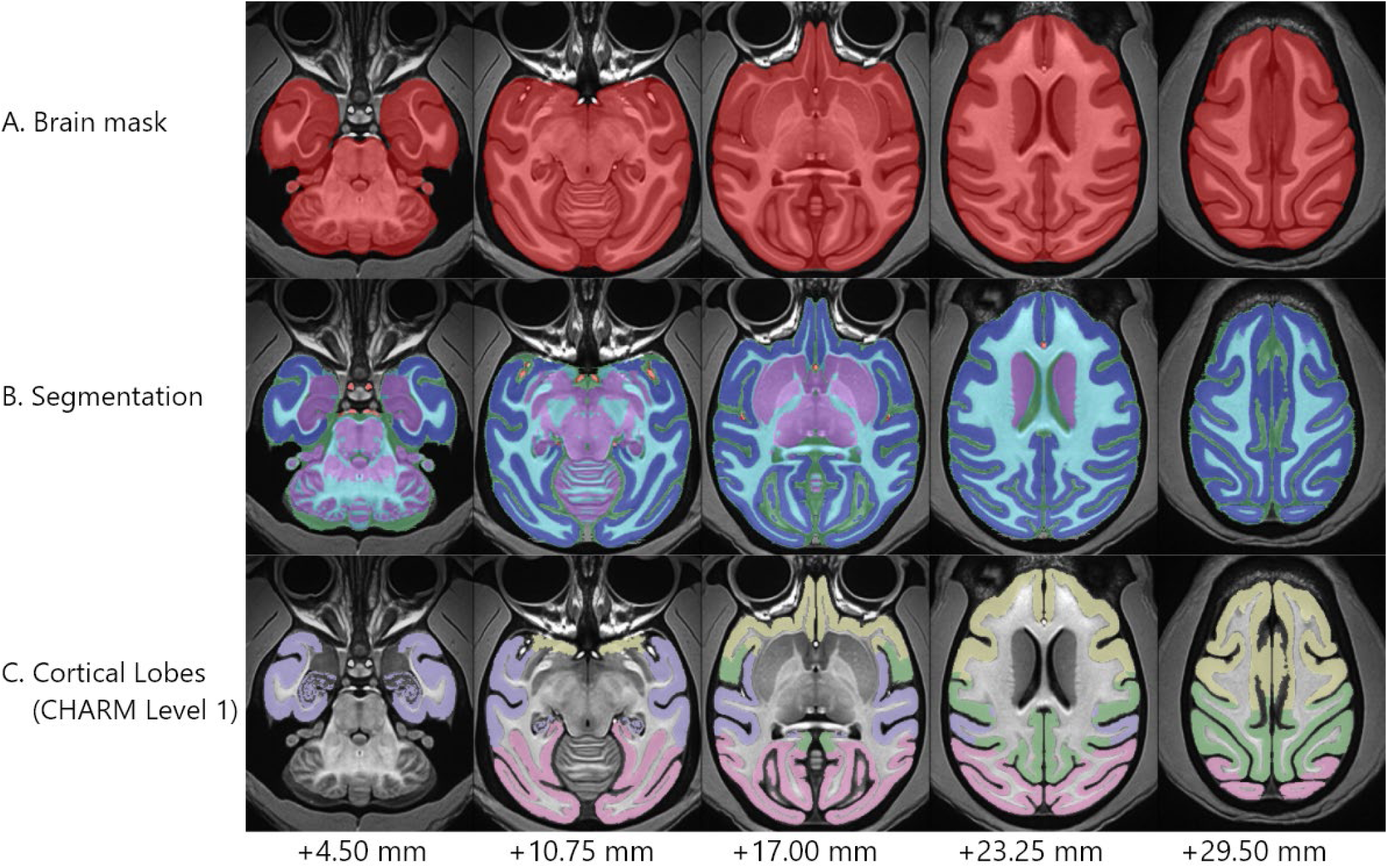
The asymmetric NMT v2 and associated datasets. Axial slices through the asymmetric NMT v2 are shown as an underlay, with various masks distributed with the template overlaid translucently. A) The asymmetric NMT v2 brain mask (red) captures the brain and associated vasculature. B) The brain mask is further divided into a 5-class segmentation of the following types: cerebrospinal fluid (CSF; green), subcortical gray matter (scGM; purple), cortical gray matter (GM; dark blue), white matter (WM; light blue) and large blood vasculature (BV; red). C) Multiple atlases are provided with the NMT v2, including the CHARM. Level 1 of the CHARM hierarchy is displayed here, which consists of the frontal (yellow), temporal (purple), parietal (green) and occipital (pink) cortical lobes. Distances are superior to the interaural meatus. The symmetric segmentation, shown in Figure 4, is quite similar.

**Figure S3.**
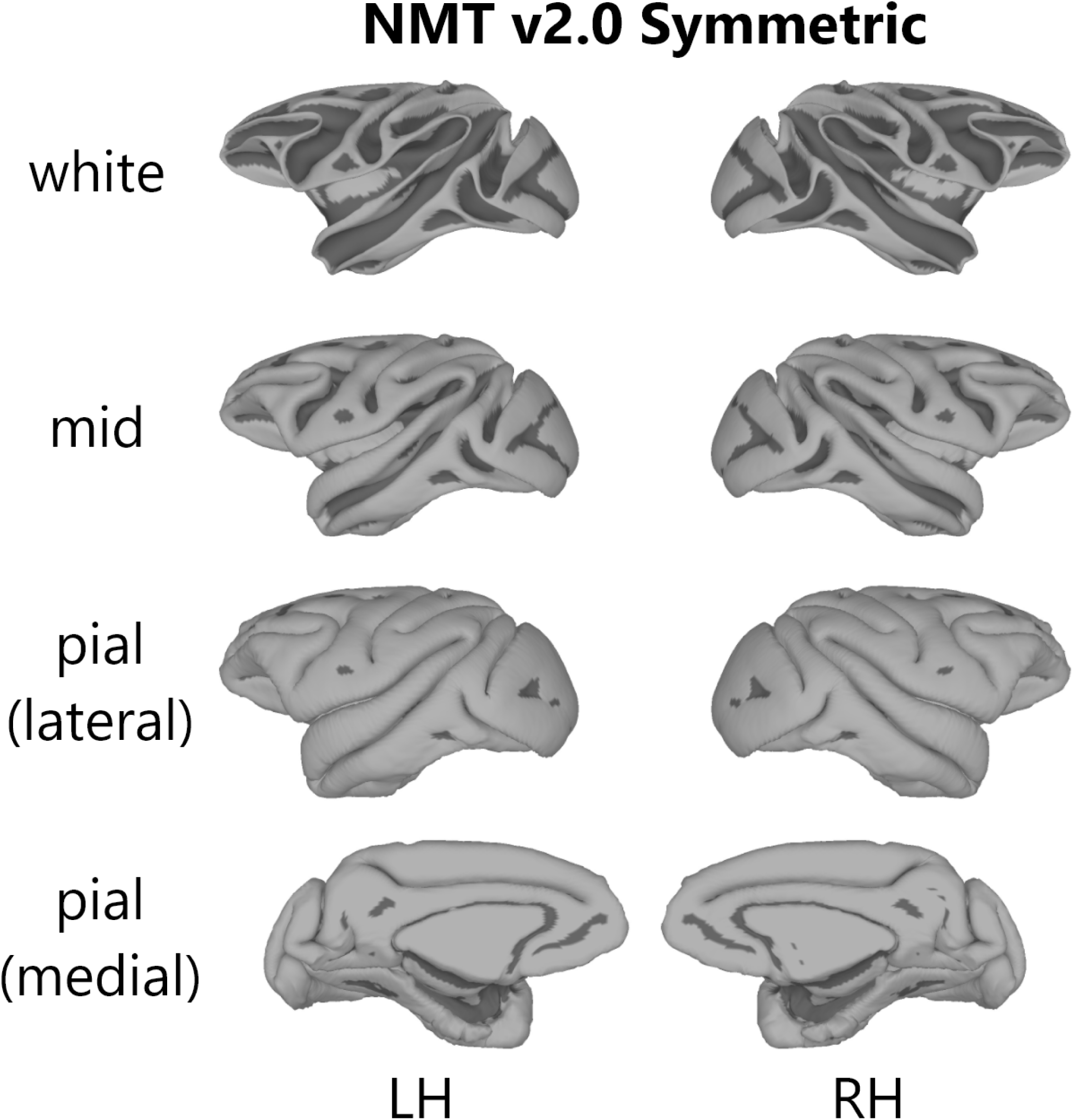
Surfaces of the symmetric NMT v2. The NMT v2 comes with white matter (i.e. the border between white and gray matter), mid-cortical (between white and pial) and pial surfaces. The right hemisphere surfaces are displayed above for the symmetric NMT v2 (both hemispheres are provided in the NMT v2). Surfaces were generated using CIVET-macaque (Lepage et al., this issue) and visualized in SUMA (Saad et al., 2004). Here, grayscale denotes convexity: light gray is positive convexity, and dark gray is negative. Semi-inflated surfaces (not shown) were also generated using the AFNI command SurfSmooth. Surfaces are available in the commonly used GIFTI format.

**Figure S4.**
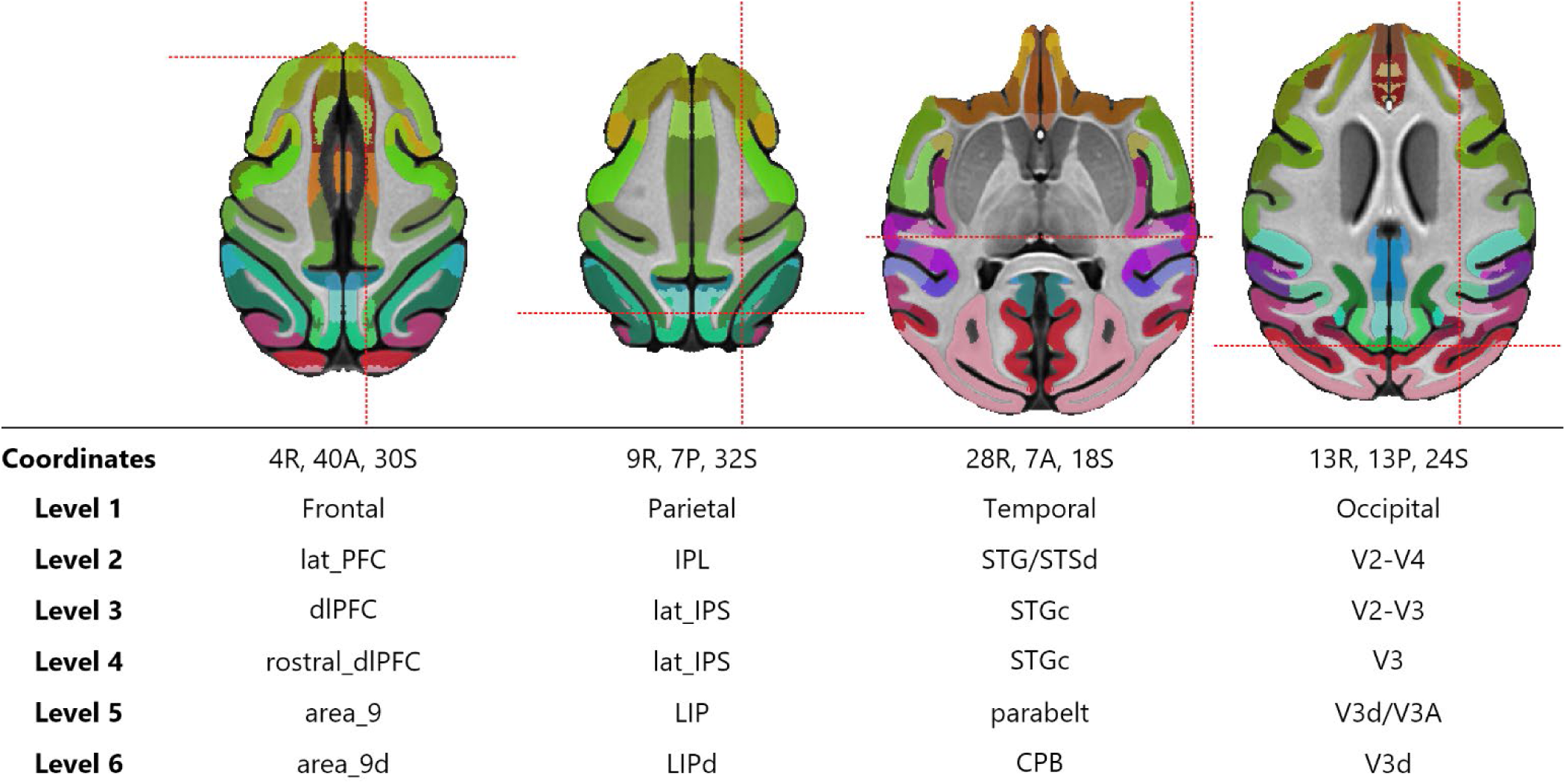
CHARM regions for an example location in each cortical lobe. Each cortical GM voxel is assigned a label at each of the 6 levels of CHARM, with some labels repeating over 2 or more levels. Shown here are 4 example voxels (red crosshair locations) with their associated ROI abbreviations across the levels of the hierarchy. Axial slices are displayed with the CHARM level 6 regions overlaid. Coordinates are in mm from the origin in the indicated direction (L/R = Left/Right, A/P = Anterior/Posterior; S/I = Superior/Inferior). Abbreviations: lat_PFC = lateral prefrontal cortex, dlPFC = dorsolateral prefrontal cortex, area_9d = area 9 [dorsal portion], IPL = inferior parietal lobule, lat_IPS = lateral intraparietal sulcus, LIP = lateral intraparietal area, LIPd = dorsal LIP, STG/STSd = superior temporal region, STSd = dorsal bank of the superior temporal sulcus, STGc = caudal superior temporal gyrus, CPB = caudal parabelt, V2-V4 = extrastriate visual areas 2-4, V2-V3 = preoccipital visual areas 2-3, V3d = dorsal visual area V3, V3A = visual area V3A.

## Footnotes

1 https://afni.nimh.nih.gov/pub/dist/doc/htmldoc/nonhuman/macaque_tempatl/template_nmtv2.html

2 https://afni.nimh.nih.gov/pub/dist/doc/htmldoc/nonhuman/macaque/demo_task_fmri.html

3 https://afni.nimh.nih.gov/pub/dist/doc/htmldoc/nonhuman/macaque/demo_rest_fmri.html

4 http://fcon_1000.projects.nitrc.org/indi/indiPRIME.html

5 In 3 subjects, we used multiple scans from different sessions. In these cases, the transformation parameters were averaged together within a subject before being averaged across subjects.

6 https://github.com/jms290/NMT

7 https://afni.nimh.nih.gov/pub/dist/doc/htmldoc/nonhuman/macaque_tempatl/template_nmtv1.html

8 http://braininfo.org/

9 https://scalablebrainatlas.incf.org/main/hierarchy.php

10 http://atlas.brain-map.org/

11 https://afni.nimh.nih.gov/pub/dist/doc/htmldoc/nonhuman/macaque_tempatl/template_nmtv2.html

12 https://afni.nimh.nih.gov/pub/dist/doc/htmldoc/index.html

13 https://github.com/afni/afni

14 https://afni.nimh.nih.gov/pub/dist/doc/htmldoc/nonhuman/macaque/demo_task_fmri.html

15 https://afni.nimh.nih.gov/pub/dist/doc/htmldoc/nonhuman/macaque/demo_rest_fmri.html

16 prime-re.github.io

17 https://afni.nimh.nih.gov/pub/dist/doc/htmldoc/nonhuman/macaque/main_toc.html

18 http://hpc.nih.gov

